# Designed high-redox potential laccases exhibit high functional diversity

**DOI:** 10.1101/2022.06.22.497129

**Authors:** Shiran Barber-Zucker, Ivan Mateljak, Moshe Goldsmith, Meital Kupervaser, Miguel Alcalde, Sarel J. Fleishman

## Abstract

White-rot fungi secrete an impressive repertoire of high-redox potential laccases (HRPLs) and peroxidases for efficient oxidation and utilization of lignin. Laccases are attractive enzymes for green-chemistry applications due to their broad substrate range and low environmental impact. Since expression of functional recombinant HRPLs is challenging, however, iterative directed evolution protocols have been applied to improve their expression, activity and stability. We implement a rational, stabilize-and-diversify strategy to two HRPLs that we could not functionally express: first, we use the PROSS stability-design algorithm to allow functional expression in yeast. Second, we use the stabilized enzymes as starting points for FuncLib active-site design to improve their activity and substrate diversity. Four of the FuncLib designed HRPLs and their PROSS progenitor exhibit substantial diversity in reactivity profiles against high-redox potential substrates, including lignin monomers. Combinations of 3-4 subtle mutations that change the polarity, solvation and sterics of the substrate-oxidation site result in orders of magnitude changes in reactivity profiles. These stable and versatile HRPLs are a step towards the generation of an effective lignin-degrading consortium of enzymes that can be secreted from yeast. More broadly, the stabilize-and-diversify strategy can be applied to other challenging enzyme families to study and expand the utility of natural enzymes.

## Introduction

Enzymatic conversion of biomass into useful materials and energy sources is critical for the drive towards a greener, more energy-efficient economy. Lignocellulose, comprising cellulose, hemicellulose and lignin, is the main component of the plant cell wall and the most abundant source of biomass on earth. The cellulose and hemicellulose components can be effectively degraded by hydrolytic enzymes or using mild chemical conditions to their monosaccharides for biofuel production^1^. Lignin, however, is a cross-linked aromatic heteropolymer that is intertwined with cellulose and hemicellulose. It is therefore extremely recalcitrant to depolymerization, requiring harsh conditions such as ionic liquids, extreme pHs or high temperatures. Economically viable and environmentally benign methods for lignin depolymerization may increase access to cellulose and hemicellulose for biofuel production and mobilize the high-value aromatic monomers stored in lignin^2–4^.

White-rot fungi secrete a broad repertoire of oxidoreductases that together with other auxiliary enzymes synergistically oxidize lignin, leading to its depolymerization^5,6^. These oxidoreductases are grouped into two major types: heme-containing peroxidases (manganese, lignin and versatile peroxidases) and copper-dependent polyphenol oxidases named laccases (E.C. 1.10.3.2). Although peroxidases have higher redox potential (up to 1.40 V vs. normal hydrogen electrode, NHE), they depend on hydrogen peroxide which leads to heme bleaching and enzyme deactivation even in moderate hydrogen peroxide concentrations^7,8^. By contrast, laccases are highly promiscuous, O_2_-dependent oxidoreductases in which paramagnetic Cu^2+^ (type 1 copper, T1Cu) oxidizes diverse substrates in a partially exposed recognition site. Following oxidation, the electrons are transferred to and stored in a buried trinuclear copper cluster (T2/T3Cu) which reduces molecular oxygen to two water molecules. Thus, laccases exhibit two key advantages for industrial application: they use molecular oxygen rather than the harsher hydrogen peroxide, and their sole byproduct is water, making them both longer lasting compared to peroxidases and milder in relation to chemical oxidation^9,10^.

Laccases from white-rot fungi are of special interest as they exhibit a higher redox potential (high redox potential laccases, or HRPLs; 0.720-0.790 V vs. NHE) compared to other fungal, bacterial or plant laccases (0.400-0.700 V vs. NHE). HRPLs do not oxidize lignin directly but through laccase mediators – small aromatic compounds that can diffuse into the laccase active site, undergo oxidation and then diffuse into the lignin mesh and attack it^9,11,12^. Despite these advantages, however, HRPLs typically express very poorly in heterologous hosts impeding research and applications. This is, among other reasons, due to their high content of irregular structures (approximately 50% of the protein comprises loops), several disulfide bonds and glycosylation sites^13,14^. Previous HRPL engineering campaigns mostly used directed evolution, including recently computation-guided evolution, HRPL chimeragenesis or *in vitro* evolution starting from a reconstructed ancestral HRPL^9,15–18^. In all these campaigns, however, the engineering of a single HRPL demands substantial and iterative experimental effort, starting from improving its heterologous expression to engineering its activity, stability and substrate specificity. Furthermore, HRPLs, peroxidases and other auxiliary enzymes attack lignin synergistically, demanding several enzymes from each family for efficient attack^5,19^. For these reasons, a common and facile expression host, such as *S. cerevisiae*, is essential for research and industrial applications, and the above-mentioned engineering difficulties present a serious bottleneck. Computational design methodologies can be used as effective alternatives to iterative engineering approaches. The PROSS^20^ and FuncLib^21^ algorithms previously developed in our lab have been used to improve diverse proteins in one-shot design; that is, without iterating modeling, mutagenesis and screening^20–25^. Both algorithms use Rosetta atomistic design calculations and phylogenetic data to restrict design to structurally tolerated substitutions that are likely to occur in natural evolution. PROSS designs mutants that exhibit higher stability and expressibility; hence, design is allowed only outside the active sites^20,22^. By contrast, FuncLib aims to improve catalytic activity or alter substrate scope, and accordingly, the design process is limited to residues in the active site^21^. Thus, PROSS and FuncLib are compatible and complementary methods that rationalize and dramatically accelerate many of the iterative steps that are necessary for enzyme optimization. We recently used PROSS to enable the functional expression of enzymes as complex as versatile peroxidases (VPs) in yeast^25^. This encouraged us to extend these methods to other ligninolytic oxidoreductases, with the ultimate aim of constructing an artificial yeast secretome for efficient lignin depolymerization^5^.

Here, we focused our design effort on three fungal HRPLs whose structures were determined but are hardly expressed functionally in yeast. We first used the PROSS stability-design calculations to enable functional expression of HRPLs from *Trametes hirsuta* and *Trametes versicolor* in yeast by introducing dozens of mutations outside the active site. We then selected the most stable and active PROSS designs for further design of the T1Cu site using the FuncLib algorithm implementing 3-7 mutations within the substrate-oxidation site. Four *Trametes hirsuta* FuncLib designs were further characterized together with their PROSS progenitor, and shown to exhibit high stability and distinct reactivity profiles against laccase substrates, including orders of magnitude differences in substrate oxidation profiles. Structural analysis reveals that the combination of subtle mutations leads to these dramatic changes in activity, demonstrating that this stabilize-and-diversify strategy can illuminate structure-function relationships even in challenging enzymes.

## Results

### Design of HRPLs for functional expression in yeast

Since many HRPLs hardly express in yeast, we began by designing more stable HRPL variants using the PROSS stability design algorithm^20^. Furthermore, activity-enhancing mutations often destabilize proteins, and a stable scaffold would permit more function-altering mutations^26^. The crystal structures of *Trametes versicolor, Trametes hirsuta* and basidiomycete PM1 HRPLs (PDB entries 1GYC^27^, 3FPX^28^ and 5ANH^29^, respectively) were used as starting points for PROSS design^20,22^ (Figure 1A,B). For each HRPL, we selected the wildtype and four PROSS designs with different mutational loads for experimental characterization (20-84 mutations in each design, corresponding to 4-16% of the protein sequence; see Table S1 and the designed sequences in SI). In three of the four designs for each wild type HRPL, we disabled mutations in known glycosylation sites and eliminated mutations that might impact the native N-glycosylation patterns, since glycosylations can enhance stability. As they are not entirely conserved in laccases, however, we further tested a single design for each HRPL which was allowed to impact glycosylation sites (designs named 9nL, where nL stands for non-limited N-glycosylations). We note that all atomistic modeling was done in the absence of glycans. The DNA encoding each protein was incorporated into the pJRoC30 plasmid and transformed into *S. cerevisiae*. Since the wild type enzymes could not be functionally expressed in this system, as a reference for stability and activity, we used the highly secreted, stable and active basidiomycete PM1 variant OB-1, which was optimized by *in vitro* evolution^30^.

**Figure 1.**
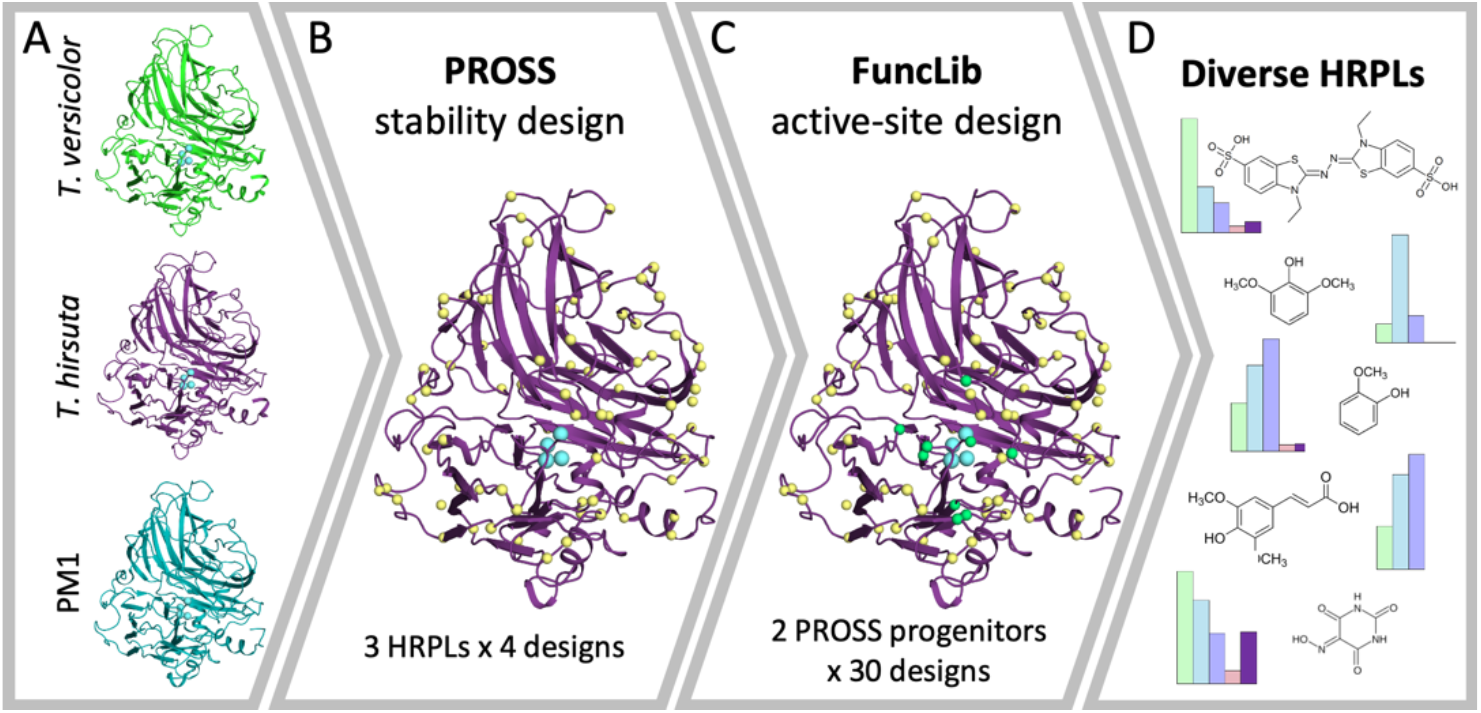
The Stabilize-and-diversify strategy. (A) The crystal structures of three HRPLs from *Trametes versicolor, Trametes hirsuta* and basidiomycete PM1 (PDB entries 1GYC^27^, 3FPX^28^ and 5ANH^29^, respectively) were selected as starting points for design. (B) First, the three HRPLs were designed for expressibility and stability using PROSS^20^. For each wild type protein, four PROSS designs were selected for experimental testing. Copper atoms (cyan) and some PROSS designed positions (yellow) are indicated in spheres. (C) Two designs with highest activity and stability, one of *Trametes versicolor* and one of *Trametes hirsuta*, were selected for active-site design (green spheres) using FuncLib^21^. For each PROSS-designed progenitor, 30 designs were selected for experimental screening. (D) FuncLib designs show diverse reactivity profiles. Four designs from *Trametes hirsuta* (light pink, light purple, light blue and light green) show dramatic improvements in catalytic efficiency against various HRPL substrates compared to their PROSS progenitor (purple).

We initially screened all PROSS designs against the general oxidoreductase substrate 2,2′-azino-bis (3-ethylbenzothiazoline-6-sulfonic acid) (ABTS). The screen indicated that while the wild type progenitors of all the three HRPLs exhibited no functional expression, three PROSS designs of both *Trametes versicolor* (Tv) and *Trametes hirsuta* (Th) could be functionally expressed in yeast (Figure S1A). Two designs based on the laccase from *Trametes versicolor* (Tv2 and Tv9nL with 28 and 79 mutations respectively) and three designs of *Trametes hirsuta* (Th3, Th7 and Th9 with 20, 35 and 60 mutations, respectively) were further characterized, revealing diverse stability and reactivity profiles (Figure 2 and Figure S1). For instance, Tv9nL with 79 mutations (representing 16% of the protein) — to our knowledge, an unprecedented mutational load for stability design — is highly thermostable, even compared to the evolved OB-1 (Figure 2A). Furthermore, Tv9nL exhibits remarkable pH stability (Figure 2B and Figure S1D), including high stability under very acidic pH (85% residual activity after one-week incubation at pH 2). Such acidic conditions typically destabilize HRPLs, albeit, paradoxically, lignin degradation is favored by high acidity^31,32^. This design is remarkable both for its unusually high mutational load and because it modifies four putative N-linked glycosylation sites: substitution His55Pro, Asn141Asp and Thr253Pro abolish three glycosylation motifs, and Ile301Asn introduces an Asn-X-Thr putative glycosylation motif. In fact, the glycosylation sites seen in the crystallographic structure of Tv at positions Asn54 and Asn251 are likely to be eliminated by these mutations. Our observation that glycosylation sites can in some cases be mutated without loss in activity or stability (and in fact, exhibiting a gain in activity by 2-5 fold relative to more restricted designs) may have implications for optimizing other glycoproteins.

**Figure 2.**
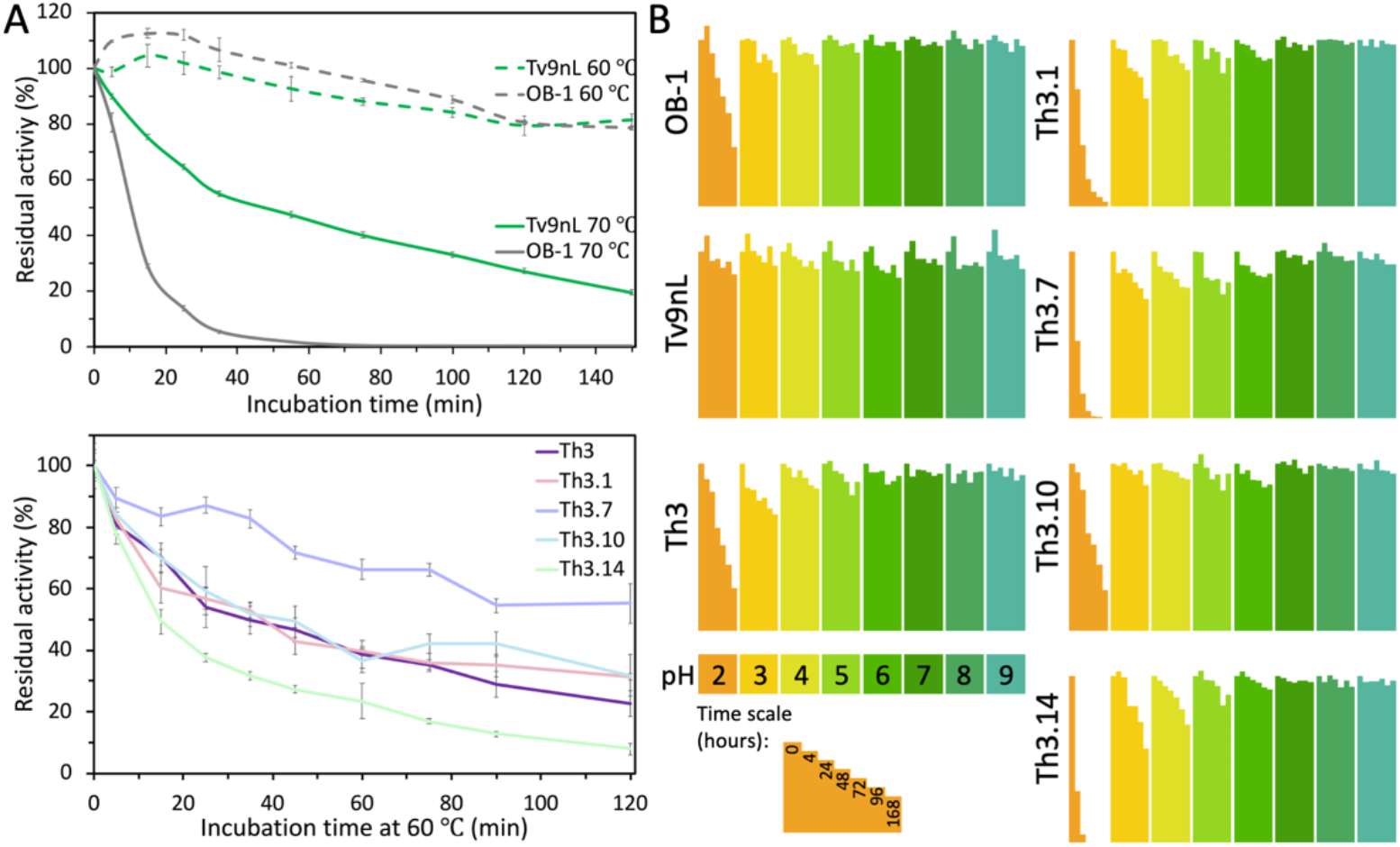
High thermal and pH stability in designs. (A) Kinetic thermostability (t_1/2_) profiles were determined by incubating yeast supernatants at 60 or 70 °C and measuring the residual activity at times 0-120 or 0-150 minutes, compared to the initial activity. (B) pH stability profiles were determined by incubating the supernatants at 100 mM borate-citrate-phosphate buffer with pH values ranging from 2 to 9 and measuring the residual activity at times 0-168 hours, compared to the initial activity at each pH. Results are presented as the mean ± S.D. of three independent experiments.

Th3, originating from a different HRPL and bearing only 20 mutations, shows low stability when compared to Tv9nL and OB-1 but high stability when compared to the other Th variants (Figure 2 and Figure S1C & D). Nevertheless, it exhibits high activity compared to all other designs, including against the HRPL high-redox potential mediator violuric acid (VLA, Figure S1A & B). Owing to their superior characteristics, Tv9nL and Th3 were selected as starting points for active-site design by FuncLib.

### Dramatic changes in stability and substrate specificity in FuncLib designs

The second step in our stabilize-and-diversify strategy was to apply FuncLib^21^ to the T1Cu site (Figure 1C), which is responsible for substrate specificity and redox potential^9,18,33^. FuncLib design was applied to 13 positions that impact HRPL activity^15,16,29,30,34–36^ in the T1Cu site of the two starting PROSS-designed HRPLs (Tv9nL and Th3), and 30 designs per progenitor were selected for experimental screening (Table S2 & S3). We first screened all the FuncLib variants expressed under restrictive growth conditions in a 96-well plate^37^ against three laccase substrates – ABTS, guaiacol (GUA, a lignin monomer) and VLA. Screening indicated that some of the FuncLib designs exhibited different specificity profiles compared to their progenitors (Figure S2A-C). Of those, 13 Th3 designs and seven Tv9nL designs were selected for a second screen against two additional lignin monomers – 2,6-dimethoxyphenol (DMP) and sinapic acid (SA). Here, several designs showed much improved activity compared to their progenitors (Figure S2D). For example, while the PROSS designs Tv9nL and Th3 exhibited no activity against SA, the FuncLib designs Tv9nL.21, Th3.7, Th3.10 and Th3.14 were active, and Tv9nL.26 could oxidize VLA while its PROSS-designed progenitor could not. These results demonstrate that FuncLib design can produce substrate preferences that are not detected in their parental proteins.

We next investigated how the active-site mutations in the most active FuncLib designs (Tv9nL designs 21 and 26, and Th3 designs 1, 7, 10, 14) impact their stability. The mutations in the Th3 FuncLib designs only slightly reduce stability, mainly in acidic conditions (for Th3 designs 1, 7 and 14, see Figure 2B and Figure S3B), and the kinetic thermostability at 60 °C is not affected for designs 1 and 10, slightly reduced for design 14 and improved for design 7 (Figure 2A). The Tv9nL FuncLib designs, however, showed much reduced thermal- and pH-stability compared to their progenitor (Figure S3), but higher activities than Tv9nL. Nevertheless, since they showed consistently lower activity than the Th3 designs and low stability, they were not further characterized.

### Designs express as a heterogeneous mix of glycosylated forms

Following the encouraging screening results, we purified Th3 and its top four FuncLib designs, Th3.1, Th3.7, Th3.10 and Th3.14, to measure their kinetic activity and specificity profiles. We purified the designs using a similar protocol as used in the *in vitro* evolution campaign of PM1 variants^15,30^ (see *Methods*) and compared their kinetics and ability to degrade the recalcitrant dye reactive black 5 (RB5) using the laccase mediator system (see below). The purified Th3 designs exhibit the expected size calculated from their sequence (∼ 56 kDa, Figure S4A). During purification, however, we found that a large fraction of the expressed designs are heavily and heterogeneously glycosylated (observed as a high molecular weight smear in an SDS-PAGE analysis; see *Methods* and Figure S4B), indicating that the designed Th3 laccases are expressed as both non-glycosylated and several heavily glycosylated forms. Hyper and differential glycosylation patterns are often observed in secreted proteins and were observed previously in HRPLs^14,38^. In fact, hyperglycosylation may underlie the high stability observed in our designs^13,14,39^.

We next applied the N-glycosidase PNGaseF to probe the source of glycosylation. Treatment of the highly glycosylated fraction resulted in the appearance of three major bands in an SDS-PAGE (Figure S4B), and proteolytic mass spectrometry showed that all the Th3 bands contained the relevant Th3 peptides. Two bands were of a larger size compared to the calculated size of the designed Th3 laccases and one of a smaller size, suggesting the designs undergo both N- and O-glycosylation and are potentially subject to proteolytic degradation. Since further purification efforts did not yield a homogeneous fraction, and as the mass spectrometry results showed that the heterogeneous, heavily glycosylated Th3 fraction also contained some *S. cerevisiae* native contaminants, we could not precisely calculate thesecretion levels of the designed Th3s. Furthermore, the characterization of the purified Th3 designs does not precisely represent the activity observed in the combined forms secreted from yeast and should be interpreted as an estimate only.

### FuncLib designs exhibit substantial diversity in reactivity

We characterized the activity profiles of the PROSS designed Th3 and its top four FuncLib designs, Th3.1, Th3.7, Th3.10 and Th3.14 (Figure 1D). In nature, lignin is decomposed under acidic conditions, and HRPL activity is acid-dependent^31,32^. The designs exhibit significantly different pH activity profiles for different substrates (Figure 3A and Figure S5). While pH 4 is optimal for Th3 relative to all substrates except ABTS (for which the activity in pH 4 is similar to the optimum activity at pH 3), for the FuncLib designs this is not always the case. For example, Th3.1 and Th3.7 are optimal at higher pH (pH 5 with GUA, and at this pH Th3.1 is also optimal with DMP and VLA).

**Figure 3.**
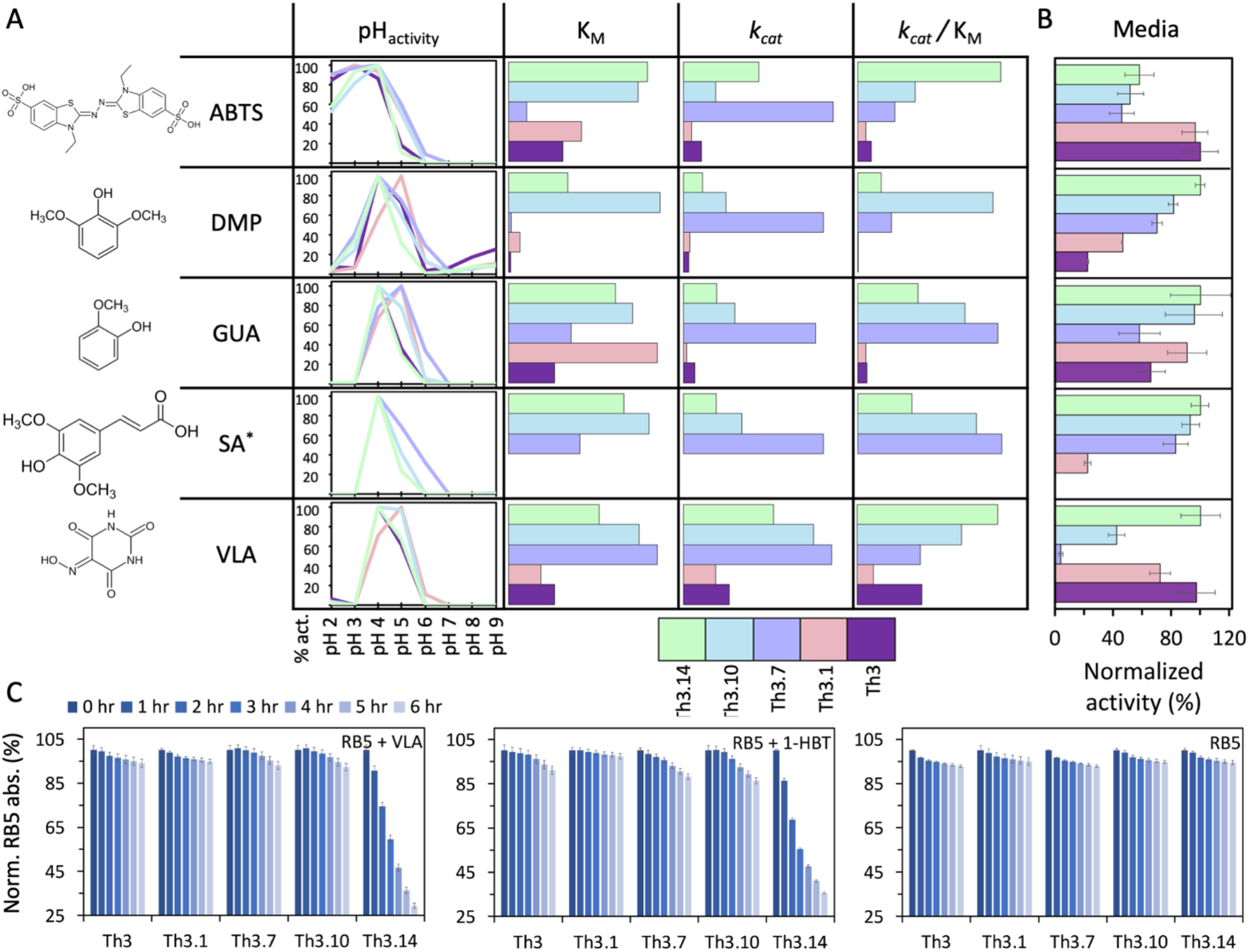
FuncLib designs exhibit striking diversity in reactivity profiles. (A) Activities of the Th3 PROSS progenitor (purple) and its four FuncLib designs (Th3.1 in light pink, Th3.7 in light purple, Th3.10 in light blue and Th3.14 in light green) were measured against five substrates (ABTS, DMP, GUA, SA and VLA). The pH-dependent activity profiles are shown as individual graphs for each substrate where the activity for each design is normalized to the activity at optimal pH. Three kinetic parameters: substrate affinity (K_M_), catalytic rate (*k*_*cat*_) or catalytic efficiency (*k*_*cat*_/K_M_) are shown. For each substrate, the bars represent the kinetic value for each design normalized to the value of the Th3 progenitor. Longer bars represent improved activities, hence lower K_M_ and greater *k*_*cat*_ and *k*_*cat*_/K_M_. For SA*, since Th3 progenitor and Th3.1 designs showed negligible activity, the values of Th3.7 and Th3.10 are similarly normalized to Th3.14. The results are presented as the mean of three independent experiments. (B) The activity of the Th3 PROSS progenitor and its four FuncLib designs was measured against the same five substrates, as in (A), but in supernatant and at saturating substrate concentrations. Here, for each substrate the activity of all designs is normalized to the activity of the design with highest activity. Results are presented as the mean ± S.D. of three independent biological replicates. (C) RB5 oxidation was determined by incubating the purified designs for six hours with RB5 and with the laccase mediators VLA and 1-HBT or without mediator and measuring the residual dye absorption at times 0-6 hours, compared to the initial absorption for each design. The results are presented as the mean ± S.D. of three independent experiments.

We further tested the Th3 designs’ activities against the five substrates at their optimal pHs. We used the purified enzymes to study their kinetics and, as these purified proteins are only one fraction of the produced enzymes, to estimate the differences between their activities. The activities of the FuncLib designs differ dramatically from those of their Th3 PROSS-designed progenitor (Figure 3A and Table S4). First, for all of the substrates, Th3 is not the best enzyme in terms of catalytic rate (*k*_*cat*_), substrate affinity (K_M_) or catalytic efficiency (*k*_*cat*_/K_M_). Second, while Th3 shows very low activity against SA (slight color developed only after a few hours of incubation), three designs gained SA oxidation activity. Third, Th3.7 exhibits the highest catalytic rate against all substrates while exhibiting the lowest affinity among the designs (except for DMP, where Th3.1 is worst and in SA where Th3.1 is inactive). This trend is expected as often high rate and substrate affinity trade off^40^, but the observed magnitude of the change for just a handful of mutations is striking. Of all the designs, Th3.1 has the lowest catalytic efficiency across the board, including when compared to Th3 (except for DMP, where its catalytic rate is slightly higher than Th3, but much lower than the other designs). Fourth, in terms of substrate affinities, the designs exhibit diversity without any of the designs standing out as the best: while Th3.14 has the greatest affinities to ABTS and VLA, Th3.10 has the greatest affinity to SA and DMP, which are chemically similar (sharing the same syringol head groups, see Figure 3), and Th3.1 has the greatest affinity to GUA. Last, all these differences reflect well in the catalytic efficiencies: Th3.10 shows remarkable improvement for DMP of approximately 600 fold compared to its progenitor, while Th3.14 and Th3.10 show approximately 10 and 15 fold improvements in catalytic efficiency for ABTS and GUA, respectively. Thus, the FuncLib designs improve the activity of their PROSS designed progenitor in all aspects we tested and against all substrates with each showing a unique reactivity profile. Of note, we could not compare the designs’ activity profiles relative to the wild type enzyme as it is not expressible in yeast.

The different activity profiles arising from the kinetic data reflect the behavior of only the purified designs and do not necessarily represent the activity of the glycosylated forms. Since the highly glycosylated fraction could not be effectively separated to its different constituents, we next probed the activity of the supernatant against saturating substrate concentrations (V_max_), at the designs’ optimum pH (Figure 3). This is particularly relevant for our yeast-secretome motivation where all enzymes would be produced in one host and used directly from the media^5^. Here, the results differ considerably from the trends seen for the purified fractions, and the fold change in activity relative to the progenitor and between designs is much smaller. The most prominent change is observed for Th3.1, which is highly active in media but shows poor activity in its purified form. As V_max_ directly relates to both enzyme concentration and catalytic rate, this can be explained by either much higher expression levels for Th3.1 compared to the other designs, or by differences in activity of the highly glycosylated forms relative to other forms. Since the different glycosylations forms preclude calculating their expression levels, the source of this difference is elusive and may require further analysis.

Last, we assessed the potential of the designs to degrade complex high-redox potential compounds through the laccase mediator system. We tested the purified enzymes’ ability to oxidize the dye RB5 (E=0.92 V vs. NHE) through two high-redox potential mediators, VLA (E=0.92 V vs. NHE) and 1-hydroxybenzotriazole (1-HBT, E=1.08 V vs. NHE)^15^. While neither Th3 nor its FuncLib designs oxidize RB5 effectively in the absence of mediator, Th3, Th3.7 and Th3.10 show improved oxidation of RB5 through both VLA and 1-HBT, and here too, the FuncLib designs are better than their progenitor. Importantly, Th3.14 efficiently oxidizes RB5 through both VLA and 1-HBT (Figure 3C) consistent with its observed high catalytic efficiency against VLA.

### Structural basis for functional diversity in designs

Lacasse structure-function studies are typically based on expert-guided substitutions in the active site^18^. Such mutations usually lead to loss-of-function, or are studied individually on a specific active-site background. Furthermore, strategies such as directed evolution only rarely mutate the active site^41^. Since each FuncLib design carries 3-4 active-site mutations from its PROSS-designed progenitor, they allow us to explore structure-activity relations in complex multipoint active-site mutants.

The T1Cu site contributes significantly to laccase redox potential^9,18,33^. For example, T1Cu shielding and hydrophobicity increase redox potential^42^, and therefore vicinal polar-to-hydrophobic mutations often result in enhanced activity and *vice versa*. By contrast with these observations, Th3.7, which has the highest catalytic rate against all substrates, carries two unique deshielding and polar-to-charged mutations in the T1Cu binding site: Ile455Val and Asn206Asp. Position 455 is in the first shell of T1Cu, eliminating a methyl group and partly deshielding T1Cu. The vicinal mutations Asn206Asp and the additional Th3.7 Ser427Asp mutation polarize the T1Cu site (Figure 4A). Nonetheless, we observed a high catalytic rate for Th3.7, demonstrating that, surprisingly, in some contexts enhanced polarity does not reduce activity. Interestingly, only one additional FuncLib design introduces the Asn206Asp and Ile455Val mutations (design Th3.8 with low activity for most substrates, see Table S3 and Figure S2D). While Th3.7 bears the additional mutation Ser427Asp, Th3.8 has the native Ser427, suggesting that this mutation compensates for the increased polarity.

**Figure 4.**
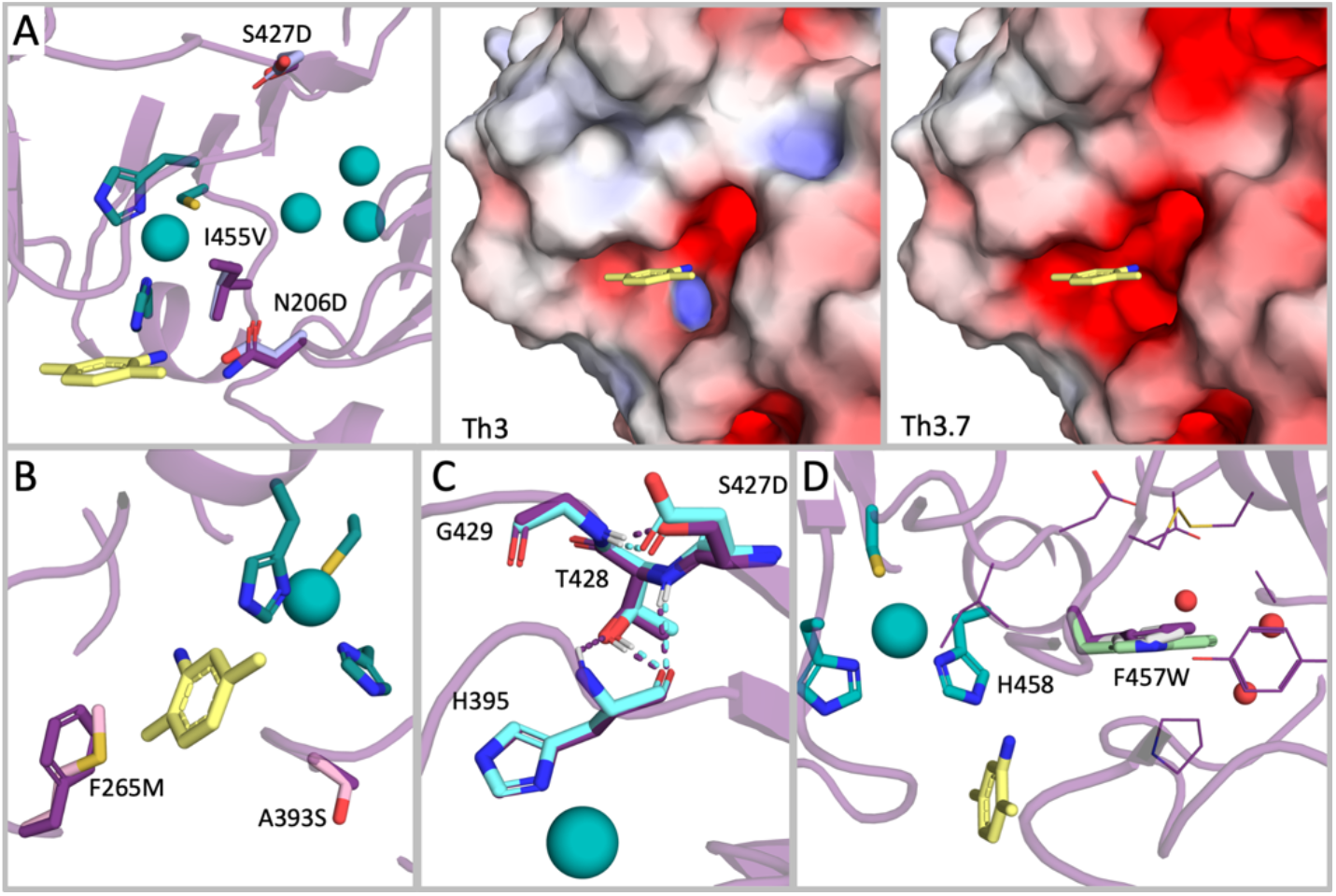
Subtle changes to active-site electrostatics, solvation and packing underlie the dramatic reactivity differences in designs. Th3 backbone, based on PDB entry 3FPX^28^, is presented as a purple cartoon. Th3 is colored in purple, Th3.1 in light pink, Th3.7 in light purple, Th3.10 in light blue and Th3.14 in light green in all panels. The four copper atoms are presented as teal spheres and T1Cu chelating residues from Th3 are presented in teal sticks. 2,5-xylidine from the *Trametes versicolor* laccase structure (PDB entry 1KYA^44^) located in the T1Cu binding site is shown in yellow sticks. (A) Changes in T1Cu hydrophobicity. Substitutions Asn206Asp, Ser427Asp and Ile455Val in Th3.7 are presented as sticks (left). Electrostatic potential maps of Th3 (middle) and Th3.7 (right) T1Cu sites were calculated using the APBS suite^45^ through the PyMOL Molecular Graphics System (Version 2.4.1, Schrödinger, LLC) and are colored in scale of [-5,5] kT/e. (B) Changes in substrate accessibility. Substitutions Phe265Met and Ala393Ser in Th3.1 are presented in sticks. (C) Changes in T1Cu-chelating His395 environment. Substitution Ser427Asp in Th3.10 and neighboring positions. Hydrogen bonds within the 427-429 triad and with His395 are indicated in dashed lines and colored as the designs. (D) Dense packing near the T1Cu site and obstruction of water molecules. Substitution Phe457Trp in T3.14 is shown in sticks, and residues in close proximity to Th3 Phe457 are presented as purple lines. Phe457 from *Trametes hirsuta* laccase crystal structure (pdb entry 3FPX^28^) is presented as gray sticks, and proximal water molecules as red spheres.

T1Cu is also the substrate recognition site, hence mutations can also change substrate preferences. Asp206 is conserved in basidiomycetes^35^ and forms critical hydrogen bonds with laccase substrates, reducing the electron transfer distance between the substrate and T1Cu-chelating His458^43^. In *Trametes hirsuta* laccase, however, position 206 exhibits an Asn. Th3.10 has the highest affinity for DMP (K_M_=17.1 and 1280 μM for Th3.7 and Th3, respectively) and SA (K_M_=150 μM for Th3.7 and Th3’s K_M_ could not be determined) which share the same 2,6-dimethoxyphenol (syringol) functional group. Here, we propose that the addition of negative charge (Asn206Glu) to the binding pocket stabilizes the substrate^35^ in Th as well. Furthermore, Th3.14, which shows the second-best affinity to these substrates (K_M_=44 and 185 μM for DMP and SA, respectively) carries the Asn206His substitution that also extends into the binding cleft but has a different impact on electrostatics. Last, Th3.7 carries Asn206Asp which changes binding-site electrostatics, while Th3 and Th3.1 bear the native Asn206 residue (Figure 4A). Th3 and Th3.1 have dramatically lower kinetic parameters towards DMP and SA (see Figure 3A & B and Table S4). Of the 30 FuncLib designs, only designs 1, 6, 25 and 26 maintain the native Asn206, and while all of them are active against ABTS, they exhibit poor activity against DMP and SA (Table S3 and Figure S2). Since 206 is the only polar position mutated in the characterized designs that faces directly into the oxidation site, our results suggest that Asn in this position reduces dramatically the activity against syringol-type substrates. This is in agreement with previous studies of fungal laccases where mutations from Asp or Glu to Asn changed dramatically the activity towards DMP^9,34,35^.

Finally, we observe that mutations in surface exposed positions in the substrate entry pathway impact activity^29,36^. Two unique mutations in surface-exposed positions in Th3.1, Phe265Met and Ala393Ser, which may interfere with the substrate entry to the T1Cu site, may underlie the comparably low activity of this design (Figure 4B). Position 427 is a second-shell position, mutations in which can impact the rigidity and stability of the T1Cu-chelating His395. The native Ser427 hydrogen bonds with Gly429, but the Ser427Asn or Ser427Asp mutations in the designs can change the hydrogen bond network with positions 428 and 429, the first of which hydrogen bonds to His395. By influencing His395’s conformation and adding polar moieties close to the T1Cu (< 10 Å), such mutations can impact electron abstraction by the T1Cu site^30^ (Figure 5C). Last, position Phe457, which is adjacent to the T1Cu-chelating position His458, is substituted for Tyr or Trp in designs Th3.1, Th3.10 and Th3.14. According to the models, these mutations improve the packing with several active-site and distant loops, potentially enhancing the rigidity of the helix on which His458 resides. Furthermore, introducing a larger side chain into a hydrophobic region may contribute to the desolvation of the T1Cu site, thus decreasing its polarity (Figure 5D). Of all the Funclib designs, only Th3.7 maintains the native Phe457, and it exhibits a shift towards activity at alkaline pHs and the highest catalytic rates. Although Phe457 does not face T1Cu (T1Cu-Phe457Cβ distance is 8 Å), mutations to polar identities at this position increase activity at alkaline pH^34^, similar to our observation with the smaller native Phe. This indicates that space-filling mutations at this position as well as its ability to rigidify His458 may dramatically impact activity and pH sensitivity. To conclude, the structure-activity analysis demonstrates that even subtle mutations (for example, eliminating a single methyl group near the copper site, or adding a hydroxyl at a second-shell position) which are unlikely to have a large impact on activity when introduced singly, can lead to dramatic changes in activity when applied simultaneously in the active site.

## Discussion

Lignin is the second most abundant terrestrial biopolymer. Due to its recalcitrance, however, it is not effectively valorized, and at present it is mostly treated as an environmental burden^4,46^. In nature, lignin is decomposed effectively only by white-rot fungi which secrete a large repertoire of oxidoreductases and auxiliary enzymes for synergistic lignin oxidation. Accordingly, adapting the fungal secretome to industrial needs is a promising avenue for environmentally benign and economically attractive lignin utilization. Because of the high complexity of the lignin substrate, however, it is essential to include in such a secretome not only one enzyme of each type, but several paralogues with different catalytic activities for each of the lignin monomers^5,19,47^. Thus, functional expression of dozens of highly stable enzymes with diverse substrate specificities in one host organism is imperative to enable efficient lignin degradation. Since the expression of the fungal enzymes in an industrially relevant organism is challenging, experimental protein engineering methods are impractical. Furthermore, these methods usually implement only very few mutations in the active site and hence are limited in the number of variants they can generate with versatile reactivity profiles^18,41^. We recently used PROSS to generate several stable and functionally diverse VPs^25^. Here, we used a different approach for diversification, demonstrating that evolution-guided atomistic design methods^48^ can be used to construct a repertoire of stable and diverse HRPLs using a two-step design protocol, again with a very low experimental effort. This stabilize-and-diversify strategy dramatically increases the success rate in creating functional diversity relative to even the most advanced laboratory engineering and evolution strategies.

Both the PROSS and FuncLib designs generated here possess attractive properties for synergistic lignin depolymerization, as well as for other industrial applications. First, the designs are stable at elevated temperatures and in diverse pH ranges. Of all the designs, the Tv9nL PROSS design stands out with high thermostability and stability at very acidic pHs which are desirable conditions for efficient lignin oxidation. Second, the designs show different pH-dependent activity profiles. For example, Th3.7 shows an activity shift to more alkaline pHs and may find uses in the many possible applications for HRPLs under physiological conditions^34^. Third, the designs show significant substrate specificity, gain activity against one substrate and reach up to 600-fold improvement in catalytic efficiency compared to their progenitor with another substrate. This indicates that the designs indeed comprise an effective repertoire of high-efficiency oxidases for different applications depending on the substrate and required reaction conditions. Moreover, the large diversity in substrate specificity profiles suggests that an enzyme cocktail exhibiting various substrate affinities and catalytic rates against each lignin monomer may act more effective on native lignin.

The stabilize-and-diversify approach is general and can be translated essentially to any enzyme family. As we show here for HRPLs, this pipeline can also highlight beneficial substitutions and serve as a platform for structure-function studies. In this case, the combination of very subtle active-site mutations yielded surprisingly large changes in activity profiles. Using FuncLib to generate a set of diverse active-site designs can therefore be a powerful strategy to extend our understanding of the rules that govern activity and substrate scope. Furthemore, the ability to heterologously express the enzymes and produce them effectively in the lab sheds light on the importance of other unanticipated molecular features, such as the existence of multiple glycosylation forms and their impact on activity. Thus, the PROSS-FuncLib combination can generate stable, diverse and efficient enzymes to address basic research and biotechnological challenges.

The mass spectrometry proteomics data have been deposited to the ProteomeXchange Consortium via the PRIDE^49^ partner repository with the dataset identifier PXD034630 and 10.6019/PXD034630.

## Acknowledgments

The authors thank Prof. Itzhak Hadar and Dr. Shira Albeck for fruitful discussion, to Prof. Avraham A. Levy, Dr. Cathy Melamed-Bessudo, Dr. Ely Morag and Dr. Rivka Elbaum for sharing their equipment and materials, and to members of the Fleishman lab for comments and advice. Research in the Fleishman lab was supported by an Individual Grant from the Israel Science Foundation (1844/19), a Consolidator Award from the European Research Council (815379), by the Sustainability and Energy Research Initiative, the Dr. Barry Sherman Institute for Medicinal Chemistry and by a donation in memory of Sam Switzer.

