## Supplemental Information for "Designed high-redox potential laccases exhibit high functional diversity"

#### Methods

**PROSS stability design calculations.** The crystal structures of *Trametes versicolor*, *Trametes hirsuta* and basidiomycete PM1 HRPLs (PDB entries 1GYC[1], 3FPX[2] and 5ANH[3], respectively) were used as starting points for PROSS design. Calculations were executed using the default web-server parameters (http://pross.weizmann.ac.il). Mutations in residues surrounding the copper atoms and at known glycosylation sites (only Asn) and cystines were disabled. For the three designs that restricted changes in glycosylation sites, mutations to and from surface Asn were disallowed prior to the PROSS design step, and further mutations that could impact glycosylations were disabled during PROSS-results analysis.

**FuncLib active-site design calculations.** The PROSS designs of Tv9nL and Th3 were used as templates for FuncLib calculations. FuncLib design was applied to residue numbers 162, 164, 206, 265, 332, 337, 393, 397, 427, 455, 457, 461, 464 in the T1Cu surrounding of both Tv9nL and Th3 (numbering relative to PDB entries 1GYC and 3FPX). Mutations to identities that are not observed frequently in evolution (PSSM score < 0) or to cysteine were eliminated (the latter, to reduce the risk of mutations that impact copper reduction). 30 designs for each progenitor were selected for experimental screening based on Rosetta energies and sequence diversity. Five designs exhibited the lowest energies and were different by at least two mutations from one another and the progenitor. The remaining 25 designs were selected by clustering all the designs with energies lower than the progenitor into 25 groups based on sequence similarity. From each cluster, the sequence that is most similar to the cluster's representative sequence (maximum one mutation from it) and has the lowest energy was selected. All the 30 selected designs differ by at least two mutations.

**Reagents.** The protease-deficient *S. cerevisiae* strain BJ5465 (α ura3-52 trp1 leu2Δ1 his3Δ200 pep4::HIS3 prb1Δ1.6R can1 GAL) was obtained from LGCPromochem (Barcelona, Spain). The uracil-independent and ampicillin resistance shuttle vector pJRoC30 was obtained from the California Institute of Technology (Caltech, Pasadena, CA). The evolved α-factor prepro-leader sequence and all the laccase genes sequences were ordered from Twist Biosciences (South San Francisco, CA). The BamHI and XhoI restriction enzymes and PNGaseF were ordered from New England Biolabs (NEB, Rehovot, Israel). ABTS, GUA, VLA, 1-HBT, RB5 and the *S. cerevisiae* transformation kit, were purchased from Sigma–Aldrich (Rehovot, Israel). DMP was purchased from Acros Organic (Geel, Belgium) and SA from Alfa Aesar (Thermo Fisher Scientific, Lancashire, UK).

**Cloning of laccases genes.** Cloning of all laccases genes was performed using the *S. cerevisiae* homologous recombination machinery[4]. pJRoC30-AAO (aryl-alcohol oxidase) expression shuttle vector previously constructed in the Alcalde lab[5] was digested with BamHI and XhoI restriction enzymes to remove the signal peptide and the AAO gene constructed within it. For PROSS designs, the evolved α-factor prepro-leader DNA sequence that was engineered in a previous directed evolution campaign of PM1L[6] (including additional restriction site in its 3' that encodes for Glu-Phe dipeptide in the N-terminal of the mature proteins; named PM1α), was ordered as a gene fragment with 40 bp overlap to the linearized plasmid. PROSS-laccase synthetic gene fragments, codon-optimized for expression in *S. cerevisiae*, were ordered each with C-terminal 6xHis-tag downstream to a thrombine cleavage site, and with 40 bp overlap to the signal peptide sequence, and to the linearized plasmid. The design of the 40 bp overlapping regions between the three fragments (plasmid, PM1α, laccase genes) allowed the recombination machinery of the protease-deficient *S. cerevisiae* strain BJ5465 to drive the fusion of the three DNA elements after transformation, and to form the pJRoC30-EvolvedSignalPeptide-LACgene expression shuttle vector. pJRoC30-OB-1 and pJRoC30-PM1L were constructed previously in the Alcalde lab[7] and the pJRoC30-SP (pJRoC30 bearing the native α-factor prepro-leader DNA sequence) in the Fleishman lab[8]. The RF method[9] was used to insert the thrombine cleavage site-6xHis-tag sequence to OB-1 plasmid (pJRoC30-OB-1-His; no change in functional expression was observed; data not shown) and to change the PM1L native α-factor prepro-leader DNA sequence to the PM1α (by applying three point mutations). Similarly, in FuncLib designs few DNA elements were used to construct the final vector by the recombination machinery of the protease-deficient *S. cerevisiae* strain BJ5465. Since FuncLib substitutions are found only in a specific region (residues 162-464), four DNA element were used for each PROSS progenitor: (1) the PM1α sequence with the DNA sequence encoding for residues 1-158 of the progenitor was ordered as a single gene fragment with 40 bp overlap to the linearized plasmid (PM1α-const1), (2) Funclib variable regions (encoding for residues 159-467) were ordered each with 40 bp overlap to the PM1α-const1 in the 5' end and to the const2-End in the 3' end, where (3) const2-End refer to a 300 bp gene fragment encoding the C-terminal of the progenitor (starting in residue 468 and including a C-terminal 6xHis-tag downstream to a thrombine cleavage site, and with 150 bp overlap to the linearized plasmid), and (4) the plasmid digested backbone above. Here as well, the design of > 40 bp overlapping regions between the four fragments (plasmid, PM1α-const1, laccase variable region, const2-End) allowed the *in vivo* recombination machinery to drive the fusion of the four DNA elements after transformation. All *S. cerevisiae*-transformed cells were plated in synthetic complete (SC) drop-out plates, and in each plate, selected colonies were picked and sequenced to verify the correct assembly and gene sequence.

**Culture media.** Minimal medium is composed of 6.7 g/L yeast nitrogen base, 1.92 g/L amino acids supplements (yeast synthetic drop-out medium supplements without uracil), 2% raffinose and 25 mg/L chloramphenicol. Laccases expression medium is composed of YP x1.11 medium (22.2 g/L bacto peptone and 11.1 g/L yeast extract), 67 mM $KH_2PO_4$ buffer at pH 6.0, 25 g/L ethanol, 22 g/L D-galactose, 2 mM $CuSO_4$ and 25 mg/L chloramphenicol. Selective expression medium (SEM) is composed of 6.7 g/L yeast nitrogen base, 1.92 g/L amino acids supplements (as above), 67 mM $KH_2PO_4$ buffer at pH 6.0, 25 g/L ethanol, 2% D-galactose, 1 mM $CuSO_4$ and 25 mg/L chloramphenicol. SC drop-out plates are composed of 6.7 g/L yeast nitrogen base, 1.92 g/L amino acids supplements (yeast synthetic drop-out medium supplements without uracil), 2% glucose, 20 g/L Bacto agar and 25 mg/L chloramphenicol.

**Screening of PROSS designs.** A colony from each *S. cerevisiae* clone containing the parental or PROSS-designed laccase genes was picked from an SC drop-out plate, inoculated in 1.5 mL minimal medium in a 14 mL culture tube, and incubated for 72 hours at 30 °C and 225 rpm. An aliquot of cells was removed and used to inoculate 3.5 mL of minimal medium in a new 14 mL culture tube to an $OD_{600nm}$ of 0.25-0.30, under the same conditions. The cells completed two growth phases (8-10 hours, reaching $OD_{600nm}$ ~ 1.0-1.5), then the expression medium (9 mL) was inoculated with 1 mL of the pre-culture in a 50 mL Falcon tube ($OD_{600nm}$ ~ 0.1). Cells were incubated for further 48 hours at 30 °C and 225 rpm and then centrifuged at 4000 g for 20 min at 4°C. The supernatants were removed into new tubes for further analysis. The expression protocol ran in triplicate, using the pJRoC30-AAO and pJRoC30-OB-1-His (referred as OB-1 in results) vectors as negative and positive controls, respectively.

An ABTS-based colorimetric assay was conducted to assess the designs' activity: 20 µL of supernatant were transferred into activity 96-well plates (Greiner Bio-One GmbH, Kremsmünster, Austria), and then, 180 µL of the reaction mixture were added to each row in the plate, and absorption at 418 nm was recorded immediately in a kinetic mode in a plate-reader at 25 °C (Citation5 or Synergy HTX plate readers, Bio-Tek, Bad Friedrichshall, Germany). The reaction mixture contained 100 mM citrate-phosphate buffer (pH 4.0) and 1 mM ABTS. All activities described in this paper were conducted using the same activity plates, in the same plate-readers and at 25 °C, unless specified otherwise. The activities were recorded for biological triplicates and averaged.

**Production of PROSS active designs.** A colony from each *S. cerevisiae* clone containing the relevant laccase gene was picked from an SC drop-out plate, inoculated in 10 mL minimal medium in a 100 mL Erlenmeyer, and incubated for 48 hours at 30 °C and 225 rpm. An aliquot of cells was removed and used to inoculate 50 mL of minimal medium in a 500 mL Erlenmeyer to an $OD_{600nm}$ of 0.25, under the same conditions. The cells completed two growth phases (8-10 hours, reaching $OD_{600nm} \sim 1$), then the expression medium (450 mL) was inoculated with 50 mL of the pre-culture in a 2 L Erlenmeyer ($OD_{600nm} \sim 0.1$). Cells were incubated for further ~60 hours at 30 °C and 225 rpm and then centrifuged at 4000 g for 20 min at 4°C. The supernatant was removed and stored at 4°C for further characterization of laccases in supernatant.

**PROSS designs activity against VLA.** Aliquots of 20 µL supernatants of selected PROSS designs were incubated with 180 µL of 20 mM VLA in 100 mM citrate-phosphate buffer (pH 4.0) in sealed 96-well plates at room temperature. Absorption at 515 nm was recorded at times 0, 3.5, 7, 27.5, 50.5 and 97.5 hours. All incubations and absorption recordings were conducted in triplicate and averaged.

**Kinetic thermostability ($t_{1/2}$).** Supernatants of the selected designs (PROSS active designs or selected FuncLib designs) at appropriate dilutions (with expression buffer, to achieve non-saturated linear response in kinetic mode measurements in activity reads) were incubated in a thermocycler (S1000™ thermocycler, Bio-Rad, Rishon LeZion, Israel) pre-heated to 60 °C or 70 °C. Aliquots of 25 or 30 µL were removed at different times, chilled out on ice for 10 min and further incubated at room temperature for at least 10 min. Activity at each time point was measured using the ABTS-based colorimetric assay described above and all activities were normalized to the activity at time 0 for residual activity calculations. All incubations and activity assays were conducted in triplicate and averaged.

**pH stability.** Supernatants of the selected designs (PROSS active designs or selected FuncLib designs) were diluted to reach a final concentration of 100 mM borate-citrate-phosphate buffer at pH ranging from 2-9. Aliquots of 20 µL were removed at different times (time 0, 4, 24, 48, 72, 96 and 168 hours) and measured in the ABTS-based colorimetric assay described above. For each pH, the activities were normalized to the activity at time 0 for residual activity calculations. All incubations and activity assays were conducted in triplicate and averaged.

**First screening of FuncLib designs.** A colony from each *S. cerevisiae* clone containing the parentals, PROSS progenitors or FuncLib-designed laccase genes was picked from an SC drop-out plate, inoculated in 200 µL SEM in 96 deep-well sealed plates, and incubated for 72 hours at 30 °C and 220 rpm in an air shaker. Open flasks filled with DW were shaken together with the plates to generate humidity and prevent evaporation of the samples in the plates. The expression protocol ran in triplicate, using pJRoC30-AAO and pJRoC30-SP vectors as negative controls, and pJRoC30-OB-1-His (referred as OB-1 in results) as a reference. The plates were centrifuged at 2500 g for 15 min at 4°C and aliquots of 20 µL supernatants were transferred from the three replicated master plates to activity plates, followed by addition of 180 µL of reaction mixtures (100 mM citrate-phosphate buffer pH=4.0 with 3 mM ABTS, 10 mM GUA or 20 mM VLA). The ABTS-, GUA- and VLA-based colorimetric assays were conducted to assess the designs activity profiles. For ABTS, the absorption at 418 nm was recorded in a kinetic mode immediately after the addition of reaction mix. For GUA, the absorption at 465 nm was recorded at times 0 and 3 hours, and for VLA, the absorption at 515 nm was recorded at times 0, 27 and 43 hours. The absorption and activities were recorded for the biological triplicates and averaged.

**Production of selected FuncLib designs.** A colony from each *S. cerevisiae* clone containing the relevant laccase gene was picked from an SC drop-out plate, inoculated in 1 mL minimal medium in a 14 mL culture tube, and incubated for 48 hours at 30 °C and 225 rpm. An aliquot of cells was removed and used to inoculate 2.5 mL of minimal medium in a new 14 mL culture tube to an $OD_{600nm}$ of 0.25-0.30, under the same conditions. The cells completed two growth phases (8-10 hours, reaching $OD_{600nm}$ ~ 1.0-1.5), then the expression medium (9 mL) was inoculated with 1 mL of the pre-culture in a 50 mL Falcon tube ($OD_{600nm}$ ~ 0.1). Cells were incubated for further 60-62 hours at 30 °C and 225 rpm and then centrifuged at 4000 g for 20 min at 4°C. The supernatant was removed and stored at 4°C for further analysis. For relevant assays, the expression protocol ran in triplicate.

**Activity assays of selected FuncLib designs.** Aliquots of 20 µL of selected FuncLib designs supernatants were transferred to activity plates, followed by addition of 180 µL of reaction mixtures (100 mM citrate-phosphate buffer pH=4.0 with 3 mM ABTS, 3 mM DMP, 10 mM GUA, 0.25 mM SA or 20 mM VLA). The ABTS-, DMP-, GUA-, SA- and VLA-based colorimetric assays were conducted to assess the selected designs activity profiles. The absorptions of ABTS, DMP, GUA, SA and VLA at 418, 469, 465, 512 and 515 nm, respectively, were recorded in a kinetic mode immediately after the addition of the reaction mixtures. The absorption and activities were recorded in triplicates and averaged.

**pH activity profiles of selected FuncLib designs.** Aliquots of 20 µL of selected FuncLib designs supernatants were transferred to activity plates. 180 µL of the reaction mixture was added to the plate, and absorption at the appropriate wavelength (substrate-dependent) was recorded immediately in a kinetic mode. The reaction mixtures contained a specific substrate in a 100 mM borate-citrate-phosphate buffer (pH 2, 3, 4, 5, 6, 7, 8, 9). The following substrate concentrations were used: ABTS 3 mM, DMP 3 mM, GUA 10 mM, SA 0.25 mM and VLA 20 mM, using the same absorption wavelengths described above. For each protein and substrate, the activities were normalized to the activity at optimal pH for residual activity calculations. Each activity assay was conducted in triplicate and averaged.

**Activity profiles of selected FuncLib designs in media at optimum pH.** Aliquots of 20 µL of selected FuncLib designs supernatants were transferred to activity plates. 180 µL of the reaction mixture was added to the plate, and absorption at the appropriate wavelength (substrate-dependent) was recorded immediately in a kinetic mode. Reaction mixtures contained 100 mM citrate-phosphate buffer at optimum pH for the design-substrate pair, with 3 mM ABTS, 3 mM DMP, 10 mM GUA, 1 mM SA or 20 mM VLA. The absorption and activities were recorded for biological triplicates and averaged.

**Laccase production for purification and characterization.** A colony from *S. cerevisiae* clone containing the laccase gene was picked from an SC drop-out plate, inoculated in 25 mL minimal medium in a 250 flask, and incubated for 48 hours at 30 °C and 225 rpm. An aliquot of cells was removed and used to inoculate 100 mL of minimal medium in a 1 L flask to an $OD_{600nm}$ of 0.25-0.30, under the same conditions. The cells completed two growth phases (8-10 hours, reaching $OD_{600nm}$ ~ 1-1.5), then the expression medium (450 mL) was inoculated with 50 mL of the pre-culture in a 2.5 L baffled flask ($OD_{600nm}$ ~ 0.1). Cells were incubated for a further ~60 hours at 30 °C and 225 rpm. Thereafter, cells were centrifuged at 6000 g for 20 min at 4°C, and the supernatant was collected and filtered with a 0.2 μm filter bottle.

**Laccase purification – ammonium sulfate (AS) precipitated fraction.** Filtrates were subjected to precipitation with AS in two steps: a first cut of 55%, followed by centrifugation and elimination of the precipitates, and a second cut of 80%. 20 mM Bis-Tris pH=6.5 (buffer A) was used to dissolve the pellet of the second cut, and the dissolved protein solution was shaken overnight at 4 °C for maximal recovery. The dissolved fraction was then centrifuged, filtrated, concentrated, and subjected to overnight dialysis against buffer A. Filtered fractions of the proteins after dialysis were uploaded onto a 5 mL HiTrap™ Q HP column (GE Healthcare Bio-Sciences AB, Uppsala, Sweden) pre-equilibrated with buffer A, through ÄKTA pure protein purification system (GE Healthcare Bio-Sciences AB). Proteins were eluted in a two-step linear gradient from 0 to 1 M NaCl, at a flow rate of 1 mL/min: the first phase of 0-50 % over 15 column volumes (CVs) and second phase of 50-100 % over 2 CVs. The fractions of the peak with the highest laccase activity were pooled, concentrated, and dialyzed against buffer A. Protein fractions were then uploaded at a flow rate of 0.4 mL/min onto a Superdex 75 Increase 10/300 GL (GE Healthcare Bio-Sciences AB) pre-equilibrated with buffer A, through the ÄKTA pure system. The fractions of the peak with the highest laccase activity were pooled and dialyzed against 20 mM citrate-phosphate pH=6.5 (buffer B). Pure protein samples were flash-frozen in liquid nitrogen and stored at -80 °C until further use. To eliminate bias in protein concentrations due to impurities in some of the laccase samples, protein concentration was determined by running the purified samples on SDS-PAGE and calculating the concentration of the relevant band only, using bovine serum albumin at different known concentrations for calibration.

**Kinetic parameters.** Steady-state kinetics were determined for the purified AS-precipitated fractions of Th3, Th3.1, Th3.7, Th3.10 and Th3.14, by measuring the activity (initial rates) in increasing concentrations of the substrate. The $K_M$ and $k_{cat}$ values for all substrates were calculated by fitting the results to the Michaelis-Menten model ($V_0 = k_{cat}[E][S] / (K_M + [S])$). 20 μL purified protein samples (diluted in buffer B to appropriate concentration, [E]<<[S]) were transferred into activity 96-well plates and then, 180 μL of the reaction mixture were added to each row in the plate, and absorption at the appropriate wavelength (substrate-dependent) was recorded immediately in a kinetic mode in a plate-reader at 25 °C. The reaction mixtures contained substrates at varying concentrations, in 100 mM borate-citrate-phosphate buffer at optimum pH. The following molar extinction coefficients were used to calculate the substrate/product concentration: ABTS, $\varepsilon_{418\ nm} = 36{,}000\ M^{-1}\ cm^{-1}$; DMP, $\varepsilon_{469\ nm} = 27{,}500\ M^{-1}\ cm^{-1}$; GUA, $\varepsilon_{465\ nm} = 12{,}100$

$M^{-1}$ $cm^{-1}$; SA, $\varepsilon_{512\ nm}$ = 14,166 $M^{-1}$ $cm^{-1}$; VLA, $\varepsilon_{515\ nm}$ = 98 $M^{-1}$ $cm^{-1}$. All activities were recorded in triplicate and the average velocities were used for the kinetic constants' calculations.

**RB5 decolorization assay.** Decolorization of RB5 by the laccase mediator system was determined for the purified AS-precipitated fractions of Th3, Th3.1, Th3.7, Th3.10 and Th3.14. 20 μL purified protein samples (diluted in buffer B to appropriate concentrations) were transferred into activity 96-well plates and then, 180 μL of the reaction mixture were added to the plate, and absorption at the 598 nm was recorded in a kinetic mode in a plate-reader at 25 °C. The final mixtures contained 20 or 100 nM purified laccase (for VLA and 1-HBT, respectively), 0.0075% RB5, 1 mM VLA or HBT (or DDW in the same volume in case where mediator was not used) and 100 mM borate-citrate-phosphate buffer pH 4.0. Each activity assay was conducted in triplicate and averaged.

**Laccase purification – AS non-precipitated fraction.** Filtrates were subjected to one-step precipitation with 100% AS, followed by centrifugation and elimination of the precipitates. Supernatants were filtered and diluted x2 to achieve a final concentration of 20 mM Bis-Tris pH=6.5 and 2 M AS (buffer C). The supernatants were uploaded onto a 5 mL HiTrap Phenyl (HS) FF column (GE Healthcare Bio-Sciences AB) pre-equilibrated with buffer C, through the ÄKTA pure system. Proteins were eluted in a three-step linear gradient from 2 to 0 M AS, at a flow rate of 1 mL/min: the first phase of 0-30 % over 2 CVs, a second phase of 30-60 % over 20 CVs and a third phase of 60-100 % over 2 CVs. The fractions of the peak with the highest laccase activity were pooled, concentrated, and dialyzed against buffer A. The purified and concentrated fractions appeared as high molecular weight smear in SDS-PAGE, treated with PNGaseF for their N-deglycosylation and then ran again on SDS-PAGE for the assessment of their glycosilation and purification level.

**Proteolytic mass-spectrometry.** Gel bands of the N-deglycosylated Th3 were excised from the gel, sliced into 1-2mm pieces and placed in a microcentrifuge tube. The bands were destained with 25mM $NH_4HCO_3$ in 50% acetonitrile (ACN) and then vacuum dried. Protein disulfide bonds were reduced by saturating the dry gel bands with 10 mM dithiothreitol in 25 mM $NH_4HCO_3$ at 56 °C for 1 hour and alkylated with 55 mM iodoacetamide in 25 mM $NH_4HCO_3$ in the dark for 45 min at room temperature. Bands were washed twice with 25 mM $NH_4HCO_3$ and twice with 25mM $NH_4HCO_3$ in 50% ACN, then vacuum dried followed by rehydrating with 500 ng trypsin in 25mM $NH_4HCO_3$ overnight at 37°C. Peptides were then extracted by addition of 50% ACN / 5% formic acid, vortexed, centrifuged, and the supernatant was collected. The digestions were stopped by the addition of trifluroacetic acid (1% final concentration).

ULC/MS grade solvents were used for all chromatographic steps. Each sample was loaded using split-less nano-Ultra Performance Liquid Chromatography (UPLC; 10 kpsi nanoAcquity; Waters, Milford, MA). The mobile phase was: (A) $H_2O$ + 0.1% formic acid and (B) acetonitrile + 0.1% formic acid. The peptides were then separated using an Aurora (75 μm internal diameter, 250 mm length, Bruker Daltonics, Billerica, MA) at 0.30 μL/min. Peptides were eluted from the column into the mass spectrometer using the following gradient: 2% to 30% (B) in 29 min, 30% to 90% (B) in 3 min, maintained at 90% for 0.5 min and then back to initial conditions. The nanoUPLC was coupled online through a Captive Spray emitter to an ion mobility-time of flight mass spectrometer (timsTOF Pro, Bruker Daltonics). Data was acquired in data dependent acquisition (DDA)-PASEF mode. Scan range was set to 100-1700 m/z and ion mobility 1/k0 ranged 0.60-1.60 $V^{\cdot}s/cm^2$.

Data was searched using Byonic v.4.3.4 search engine (ProteinMetrics) against the Uniprot *S. cerevisiae* proteome database (January 2022 version, 6,050 entries), concatenated with the Th3 laccase sequence and common lab contaminants. Search allowed for fixed Carbamidomethylation on C, variable oxidation on MP and variable deamidation on NQ. Results were filtered for an estimated 1% FDR on the protein level.

The mass spectrometry proteomics data have been deposited to the ProteomeXchange Consortium via the PRIDE[10] partner repository with the dataset identifier PXD034630 and 10.6019/PXD034630.

**Amino acid sequences of WT proteins and PROSS designs**

**>TvWT**

AIGPAASLVVANAPVSPDGFLRDAIVVNGVFPSPLITGKKGDRFQLNVVDTLTNHTMLKSTSIHWHGFFQAGTNWAD
GPAFVNQCPIASGHSFLYDFHVPDQAGTFWYHSHLSTQYCDGLRGPFVVYDPKDPHASRYDVDNESTVITLTDWYHT
AARLGPRFPLGADATLINGLGRSASTPTAALAVINVQHGKRYRFRLVSISCDPNYTFSIDGHNLTVIEVDGINSQPL
LVDSIQIFAAQRYSFVLNANQTVGNYWIRANPNFGTVGFAGGINSAILRYQGAPVAEPTTTQTTSVIPLIETNLHPL
ARMPVPGSPTPGGVDKALNLAFNFNGTNFFINNASFTPPTVPVLLQILSGAQTAQDLLPAGSVYPLPAHSTIEITLP
ATALAPGAPHPFHLHGHAFAVVRSAGSTTYNYNDPIFRDVVSTGTPAAGDNVTIRFQTDNPGPWFLHCHIDFHLEAG
FAIVFAEDVADVKAANPVPKAWSDLCPIYDGLSEANQ

**>Tv2**

AIGPVATLVIANAPVSPDGFLRDAIVVNGVFPGPLITGKKGDRFQLNVVNQLTNPTMLKSTSIHWHGFFQKGTNWAD
GPAFVNQCPIAPGHSFLYDFHVPDQAGTFWYHSHLSTQYCDGLRGPFVVYDPHDPHASLYDVDNESTVITLTDWYHT
AARLGPRFPLGADATLINGLGRSASTPTAPLAVINVQPGKRYRFRLISISCDANYTFSIDGHNLTVIEVDGINTQPL
LVDSIQIFAAQRYSFVLNANQPVGNYWIRANPNFGTVGFAGGINSAILRYQGAPVAEPTTTQTTSVIPLIETNLHPL
EPMPVPGSPTPGGVDYALNLAFNFNGTNFFINNASFTPPTVPVLLQILSGAQTAQDLLPAGSVYPLPPHKTIEITLP
ATALAPGAPHPFHLHGHAFAVVRSAGSTTYNYNDPIWRDVVSTGTPAAGDNVTIRFVTDNPGPWFFHCHIDWHLEAG
FAIVFAEDVADVKAANPVPKAWSDLCPIYDALPEANQ

**>Tv6**

AIGPVATLVISNAPVAPDGFTRDAIVVNGVFPGPLITGKKGDRFQLNVVNQLTNPTMLKSTSIHWHGFFQHGTNWAD
GPAFVTQCPIAPGHSFLYDFHVPDQAGTFWYHSHLSTQYCDGLRGPFVVYDPHDPHASLYDVDNESTVITLADWYHT
AAQLGPRFPLGADATLINGLGRSASTPTAPLAVINVQPGKRYRFRLISLSCDANYTFSIDGHNMTIIEVDGINTQPL
TVDSIQIFAAQRYSFILNANQPVGNYWIRANPNFGTTGFDGGINSAILRYQGAPDAEPTTTQTTSVIPLIETNLHPL
EPMPVPGEPTPGGVDYALNLAFNFNGTNFFINNASFVPPTVPVLLQILSGAQTAQDLLPAGSVYPLPPHKVIEITLP
ATAAAPGAPHPFHLHGHVFAVVRSAGSTTYNYNDPIWRDVVSTGTPAAGDNVTIRFVTDNPGPWFFHCHIDWHLEAG
FAIVFAEDVDDVKAANPVPKAWSDLCDIYDALPPANQ

**>Tv9**

AIGPVATLVITNAQVAPDGFTRDAIVVNGQFPGPLITGYKGDRFQLNVINQLTNPTMLKSTSIHWHGFFQHGTNWAD
GPAFVTQCPIAPGHSFLYDFTVPDQAGTFWYHSHLSTQYCDGLRGPFVVYDPHDPHAHLYDVDNESTVITLADWYHV
AAKLGPRFPLGADATLINGLGRSSSTPTAPLAVINVQPGKRYRFRLISISCDANYTFSIDGHNMTIIEVDGINTQPY
TVDSIQIFAAQRYSFILNANQPVGNYWIRANPNFGTTGFDGGINSAILRYDGAPDAEPTTTQTTSTIPLVETNLHPL
EPMPVPGEPTPGGVDYALNLDFNFNGTNFFINNATFVPPSVPVLLQILSGAQTAQDLLPSGSVYPLPPHKVIEITFP
ATSAAPGAPHPFHLHGHQFAVVRSAGSTTYNYNDPIWRDVVSTGTPAAGDNVTIRFVTDNPGPWFLHCHIDWHLEAG
FAIVFAEDIDDVKAANPVPQAWKDLCDIYDALPPANQ

**>Tv9nL**

AIGPVATLHITNAQVAPDGFTRDAIVVNGQFPGPLITGNKGDRFQLNVINQLTNPTMLKSTSIHWHGFFQHGTNWAD
GPAFVTQCPIAPGHSFLYDFTVPDQAGTFWYHSHLSTQYCDGLRGPFVVYDPNDPHAHLYDVDDESTVITLADWYHV
AARLGPRFPLGADATLINGLGRSSSTPTAPLAVINVQPGKRYRFRLISISCDANYTFSIDGHNMTIIEVDGIPTQPY
TVDSIQIFAAQRYSFILNANQPVGNYWIRANPNFGTTGFDGGINSAILRYEGAPDAEPTTTQTTSTIPLNETNLHPL
EPMPVPGEPTPGGVDYALNLDFNFNGTNFFINNATFVPPSVPVLLQILSGAQTAQDLLPSGSVYPLPPNKVIEITFP
ATSNAPGAPHPFHLHGHQFAVVRSAGSTEYNYDNPIWRDVVSTGTPANGDNVTIRFRTDNPGPWFLHCHIDWHLEAG
FAIVFAEDIDNVKSANPVPQAWKDLCDIYNALPPANQ

**>ThWT**

AVGPVADLTITDAAVSPDGFSRQAVVVNGVTPGPLVAGNIGDRFQLNVIDNLTNHTMLKSTSIHWHGFFQHGTNWAD
GPAFINQCPISPGHSFLYDFQVPDQAGTFWYHSHLSTQYCDGLRGPFVVYDPNDPHASRYDVDNDDTVITLADWYHT
AAKLGPRFPGGADATLINGKGRAPSDSVAELSVIKVTKGKRYRFRLVSLSCNPNHTFSIDGHNLTIIEVDSVNSQPL
EVDSIQIFAAQRYSFVLDANQAVDNYWIRANPNFGNVGFDGGINSAILRYDGAPAVEPTTNQTTSVKPLNEVDLHPL
VSTPVPGSPSSGGVDKAINMAFNFNGSNFFINGASFVPPTVPVLLQILSGAQTAQDLLPSGSVYVLPSNASIEISFP
ATAAAPGAPHPFHLHGHTFAVVRSAGSTVYNYDNPIFRDVVSTGTPAAGDNVTIRFDTNNPGPWFLHCHIDFHLEGG
FAVVMAEDTPDVKAVNPVPQAWSDLCPTYDALDPNDQ

**>Th3**

AVGPVADLTITDAVVSPDGFSRQAVVVNGVTPGPLIAGNKGDRFQLNVINNLTNHTMLKSTSIHWHGFFQHGTNWAD
GPAFINQCPIAPGHSFLYDFQVPDQAGTFWYHSHLSTQYCDGLRGPFVVYDPNDPHASMYDVDNDSTVITLADWYHT
AAKLGPRFPPGADATLINGKGRAPSDPTAELSVIKVTKGKRYRFRLVSLSCNPNFTFSIDGHNLTIIEVDSVNVQPL
EVDSIQIFAAQRYSFVLDANQPVDNYWIRANPNFGNVGFDGGINSAILRYDGAPAVEPTTNQTTSVKPLNETDLHPL
VPTPVPGSPSPGGVDKAINMAFNFNGSNFFINGVSFVPPTVPVLLQILSGAQTAQDLLPSGSVYVLPSNATIEISFP
ATAAAPGAPHPFHLHGHTFAVVRSAGSTVYNYDNPIFRDVVSTGTPAAGDNVTIRFDTNNPGPWFLHCHIDFHLEAG
FAVVMAEDLPDVKAVNPVPQAWSDLCPTYDALDPNDQ

**>Th7**

AVGPVTDLYITDAVVAPDGFSRSAVVVNGQVPGPLIVGNKGDRFQLNVINNLTNTTMLRSTSIHWHGFFQHGTNWAD
GPAFVTQCPIPPGHSFTYDFQVPDQAGTFWYHSHLSTQYCDGLRGPFVVYDPNDPHASLYDVDNDSTVITLADWYHT
AAKLGPRFPPGADATLINGKGRAPSDPTAELSVIKVTPGKRYRFRLISLSCNPNFTFSIDGHNMTIIEVDSVNVQPL
EVDSIQIFAAQRYSFILDANQPVDNYWIRANPNFGNTGFDGGINSAILRYDGAPPVEPTTNQTTSTKPLNETDLHPL
VPTPVPGQPTPGGVDKAINMAFNFNGTNFFINGVSFVPPTVPVLLQILSGAQTAQDLLPSGSVYVLPPNAVIEISFP
ATAAAPGAPHPFHLHGHTFAVVRSAGSTVYNYDNPIFRDVVSTGTPAAGDNVTIRFVTNNPGPWFLHCHIDFHLEAG
FAVVMAEDLPRVKAVNPVPQAWSDLCPTYDALDPNDQ

**>Th9**

AVGPVTDLYITDADVAPDGFSRSAVVVNGQVPGPLIVGNKGDRFQINVINNLTNTTMLRSTSIHWHGFFQHGTNWAD
GPAFVTQCPIPPGHSFTYDFTVPDQAGTFWYHSHLSTQYCDGLRGPFVVYDPNDPHKHLYDVDNDSTVITLADWYHV
AAKLGPRFPPGADATLINGLGRSPSDPTAELAVIKVTPGKRYRFRLISLSCNPNFTFSIDGHNMTIIEVDSVNVQPL
EVDSIQIFAAQRYSFILDANQPVDNYWIRANPNFGNTGFDGGINSAILRYDGAPPVEPTTNQTTSTKPLNETDLHPL
VPTPVPGQPTPGGVDLAINMAFNFNGTNFFINGVTFVPPTVPVLLQILSGAQTAQDLLPSGSVYVLPPNAVIEISFP
ATSAAPGAPHPFHLHGHTFAVIRSAGSTVYNYDNPIFRDVVSTGTPAAGDNVTIRFVTNNPGPWFLHCHIDWHLEAG
FAVVMAEDLPDVKSVNPVPQAWEDLCPIYDALDPNDQ

**>Th9nL**

AVGPVTDLYITNADVAPDGFSRSAVVVNGQVPGPLITGNKGDRFQINVINQLTNTTMLRSTSIHWHGFFQHGTNWAD
GPAFVTQCPIPPGHSFTYDFTVPDQAGTFWYHSHLSTQYCDGLRGPFVVYDPHDPHAHLYDVDDDSTVITLADWYHV
AAKLGPRFPPGADATLINGLGRSPSNPTAELAVINVTPGKRYRFRLISLSCNANFTFSIDGHNMTIIEVDGVNTQPL
EVDSIQIFAAQRYSFILDANQPVDNYWIRANPNFGNTGFDGGINSAILRYDGAPPVEPTTTQTTSTNPLNETNLHPL
VPTPVPGQPTPGGVDLAINMNFNFNGTNFFINGVTFVPPTVPVLLQILSGAQTAQDLLPSGSVYVLPPNAVIEISFP
ATSNAPGAPHPFHLHGHTFAVIRSAGSTVYNYDNPIFRDVVSTGTPANGDNVTIRFTTDNPGPWFLHCHIDWHLEAG
FAVVMAEDLPNVKAVNPVPQAWEDLCPIYNALDPNDQ

**>PM1WT**

SIGPVADLTISNGAVSPDGFSRQAILVNDVFPSPLITGNKGDRFQLNVIDNMTNHTMLKSTSIHWHGFFQHGTNWAD
GPAFVNQCPISTGHAFLYDFQVPDQAGTFWYHSHLSTQYCDGLRGPIVVYDPQDPHKSLYDVDDDSTVITLADWYHL
AAKVGPAVPTADATLINGLGRSINTLNADLAVITVTKGKRYRFRLVSLSCDPNHTFSIDGHSLTVIEADSVNLKPQT
VDSIQIFAAQRYSFVLNADQDVDNYWIRALPNSGTRNFDGGVNSAILRYDGAAPVEPTTTQTPSTQPLVESALTTLE
GTAAPGNPTPGGVDLALNMAFGFAGGRFTINGASFTPPTVPVLLQILSGAQSAQDLLPSGSVYSLPANADIEISLPA
TSAAPGFPHPFHLHGHTFAVVRSAGSSTYNYANPVYRDVVSTGSPGDNVTIRFRTDNPGPWFLHCHIDFHLEAGFAV
VMAEDIPDVAATNPVPQAWSDLCPTYDALSPDDQ

**>PM13**

SIGPVADLTISNGYVSPDGFSRQAILVNGVFPGPLITGNKGDRFQLNVINNLTNHTMLKSTSIHWHGFFQHGTNWAD
GPAFVNQCPIAPGHSFLYDFQVPDQAGTFWYHSHLSTQYCDGLRGPLVVYDPHDPHKSLYDVDDDSTVITLADWYHL
PAKVGPAVPTADATLINGLGRSPNTPNADLAVITVTKGKRYRFRLVSLSCDANYTFSIDGHSMTVIEADSVNLKPLT
VDSIQIFAAQRYSFVLNADQPVDNYWIRALPNLGTRNFDGGVNSAILRYDGAAPVEPTTTQTPSTQPLVESDLHPLE
DTAAPGNPTPGGVDYALNMAFGFAGGRFTINGVSFTPPTVPVLLQILSGAQSAQDLLPSGSVYSLPPNAVIEISMPA
TSAAPGFPHPFHLHGHTFAVVRSAGSSTYNYDNPVYRDVVSTGSPGDNVTIRFRTDNPGPWFLHCHIDWHLEAGFAV
VMAEDIQDVAATNPVPQAWSDLCATYDALSPDDQ

**>PM16**

SIGPVADLTISNGDVAPDGFSRSAILVNGVFPGPLITGNKGDRFQLNVINNLTNHTMLKSTSIHWHGFFQHGTNWAD
GPAFVNQCPIPPGHSFLYDFTVPDQAGTFWYHSHLSTQYCDGLRGPLVVYDPHDPHKSLYDVDDDSTVITLADWYHL
AAKQGPAVPTADATLINGLGRSPNTPNAPLAVITVQQGKRYRFRLVSLSCDANYTFSIDGHSMTVIEADSVNLKPLT
VDSIQIFAAQRYSFVLNADQPVDNYWIRALPNLGTRNFDGGVNSAILRYDGAPPVEPTTTQTPSTQPLVESDLHPLE
DRPAPGNPTPGGVDYALNLQFGFAGGRFTINGVSFTPPTVPVLLQILSGAQSAQDLLPSGSVYSLPPNAVIEISFPA
TSAAPGFPHPFHLHGHTFAVVRSAGSSTYNYDNPVYRDVVSTGSPGDNVTIRFRTDNPGPWFLHCHIDWHLEAGFAV
VMAEDIEDVAATNPVPQAWKDLCAIYDALSPDDQ

**>PM19**

SIGPVADLVISNGQVAPDGFTRDAILVNGTFPGPLITGNKGDRFQLNVINNLTNHTMLKSTSIHWHGFFQHGTNWAD
GPAFVNQCPIPPGHSFLYDFTVPDQAGTFWYHSHLSTQYCDGLRGPMVIYDPHDPHKHLYDVDDDSTVITLADWYHT
AAKQGPAVPTADATLINGLGRSPNTPNAPLAVITVQQGKRYRFRLISLSCDANYTFSIDGHTMTVIEVDSVNIQPLT
VDSIQIFAAQRYSFILNADQPVDNYWIRALPNLGTTNFDGGVNSAILRYDGAPPVEPTTTQTPSTHPLVESDLHPLE
DRPAPGNPHPGGVDYALNLQFGFDGGRFTINGVSFTPPTVPVLLQILSGAQTAQDLLPKGSVYTLPPNAVIEITFPA
TSAAPGFPHPFHLHGHTFAVVRSAGSSTYNYDNPIWRDVVSTGTPGDNVTIRFRTDNPGPWFLHCHIDWHLEAGFAI
VMAEDIQSVAAANPVPQAWKDLCAIYDALSPDDQ

**>PM19nL**

SIGPVADLVISNGDVAPDGFSRSAILVNGQFPGPLITGNKGDRFQLNVINQLTNHTMLKSTSIHWHGFFQHGTNWAD
GPAFVNQCPIPPGHSFLYDFTVPDQAGTFWYHSHLSTQYCDGLRGPMVIYDPNDPHKHLYDVDDDSTVITLADWYHT
AAKQGPAVPTPDATLINGLGRSPNTPTAPLAVINVQQGKRYRFRLISLSCDANYTFSIDGHNMTVIEVDGVNIQPVT
VDSIQIFAAQRYSFILNANQPVDNYWIRANPNTGTTNFDGGVNSAILRYDGAPPVEPTTTQTPSTNPLVESDLHPLE
NRPAPGEPHPGGVDYALNLNFGFNGGRFFINGVSFTPPTVPVLLQILSGAQTAQDLLPKGSVYTLPPNAVIEITFPA
TSNAPGFPHPFHLHGHTFAVVRSAGSSTYNYDNPIWRDVVSTGTPGDNVTIRFRTDNPGPWFLHCHIDWHLEAGFAI
VMAEDIENVAAANPVPQAWKDLCAIYNALSPDDQ

**Amino acid sequences of characterized FuncLib designs**

**>Th3.1**

AVGPVADLTITDAVVSPDGFSRQAVVVNGVTPGPLIAGNKGDRFQLNVINNLTNHTMLKSTSIHWHGFFQHGTNWAD
GPAFINQCPIAPGHSFLYDFQVPDQAGTFWYHSHLSTQYCDGLRGPFVVYDPNDPHASMYDVDNDSTVITLADWYHT
AAKLGPRFPPGADATLINGKGRAPSDPTAELSVIKVTKGKRYRFRLVSLSCNPNFTFSIDGHNLTIIEVDSVNVQPL
EVDSIQIFAAQRYSFVLDANQPVDNYWIRANPNMGNVGFDGGINSAILRYDGAPAVEPTTNQTTSVKPLNETDLHPL
VPTPVPGSPSPGGVDKAINMAFNFNGSNFFINGVSFVPPTVPVLLQILSGAQTAQDLLPSGSVYVLPSNATIEISFP
ATAAAPGSPHPFHLHGHTFAVVRSAGSTVYNYDNPIFRDVVNTGTPAAGDNVTIRFDTNNPGPWFLHCHIDWHLEAG
FAVVMAEDLPDVKAVNPVPQAWSDLCPTYDALDPNDQ

**>Th3.7**

AVGPVADLTITDAVVSPDGFSRQAVVVNGVTPGPLIAGNKGDRFQLNVINNLTNHTMLKSTSIHWHGFFQHGTNWAD
GPAFINQCPIAPGHSFLYDFQVPDQAGTFWYHSHLSTQYCDGLRGPFVVYDPNDPHASMYDVDNDSTVITLADWYHT
AAKLGPRFPPGADATLINGKGRAPSDPTAELSVIKVTKGKRYRFRLVSLSCDPNFTFSIDGHNLTIIEVDSVNVQPL
EVDSIQIFAAQRYSFVLDANQPVDNYWIRANPNFGNVGFDGGINSAILRYDGAPAVEPTTNQTTSVKPLNETDLHPL
VPTPVPGSPSPGGVDKAINMAFNFNGSNFFINGVSFVPPTVPVLLQILSGAQTAQDLLPSGSVYVLPSNATIEISFP
ATAAAPGAPHPFHLHGHTFAVVRSAGSTVYNYDNPIFRDVVDTGTPAAGDNVTIRFDTNNPGPWFLHCHVDFHLEAG
FAVVMAEDLPDVKAVNPVPQAWSDLCPTYDALDPNDQ

**>Th3.10**

AVGPVADLTITDAVVSPDGFSRQAVVVNGVTPGPLIAGNKGDRFQLNVINNLTNHTMLKSTSIHWHGFFQHGTNWAD
GPAFINQCPIAPGHSFLYDFQVPDQAGTFWYHSHLSTQYCDGLRGPFVVYDPNDPHASMYDVDNDSTVITLADWYHT
AAKLGPRFPPGADATLINGKGRAPSDPTAELSVIKVTKGKRYRFRLVSLSCEPNFTFSIDGHNLTIIEVDSVNVQPL
EVDSIQIFAAQRYSFVLDANQPVDNYWIRANPNFGNVGFDGGINSAILRYDGAPAVEPTTNQTTSVKPLNETDLHPL
VPTPVPGSPSPGGVDKAINMAFNFNGSNFFINGVSFVPPTVPVLLQILSGAQTAQDLLPSGSVYVLPSNATIEISFP
ATAAAPGAPHPFHLHGHTFAVVRSAGSTVYNYDNPIFRDVVDTGTPAAGDNVTIRFDTNNPGPWFLHCHIDYHLEAG
FAVVMAEDLPDVKAVNPVPQAWSDLCPTYDALDPNDQ

**>Th3.14**

AVGPVADLTITDAVVSPDGFSRQAVVVNGVTPGPLIAGNKGDRFQLNVINNLTNHTMLKSTSIHWHGFFQHGTNWAD
GPAFINQCPIAPGHSFLYDFQVPDQAGTFWYHSHLSTQYCDGLRGPFVVYDPNDPHASMYDVDNDSTVITLADWYHT
AAKLGPRFPPGADATLINGKGRAPSDPTAELSVIKVTKGKRYRFRLVSLSCHPNFTFSIDGHNLTIIEVDSVNVQPL
EVDSIQIFAAQRYSFVLDANQPVDNYWIRANPNFGNVGFDGGINSAILRYDGAPAVEPTTNQTTSVKPLNETDLHPL
VPTPVPGSPSPGGVDKAINMAFNFNGSNFFINGVSFVPPTVPVLLQILSGAQTAQDLLPSGSVYVLPSNATIEISFP
ATAAAPGAPHPFHLHGHTFAVVRSAGSTVYNYDNPIFRDVVNTGTPAAGDNVTIRFDTNNPGPWFLHCHIDWHLEAG
FSVVMAEDLPDVKAVNPVPQAWSDLCPTYDALDPNDQ

All sequences N-terminus (PM1α):

MRFPSIFTADLFAASSALAAPVKTTTEDETAQIPAEAVIGYSDLEGDFDVAVLPFSNSTNNGLLFINTTIASIAAKE
EGVSLEKRETEAEF

All sequences C-terminus (thrombin cleavage site – 6xHis tag):

SSGLVPRGSSAHHHHHH

#### Supplementary Tables

**Table S1.** Laccase origins, PDB templates, protein lengths and number of mutations in each design

| | Specie | PDB entry | Protein length | # Mutations | % Identity, WT |
|---|---|---|---|---|---|
| TvWT | *Trametes versicolor* | 1GYC | 499 | 0 | 100 |
| Tv2 | | | | 28 | 94.4 |
| Tv6 | | | | 50 | 90.0 |
| Tv9 | | | | 69 | 86.2 |
| Tv9nL | | | | 79 | 84.2 |
| ThWT | *Trametes hirsuta* | 3FPX | 499 | 0 | 100 |
| Th3 | | | | 20 | 96.0 |
| Th7 | | | | 35 | 91.0 |
| Th9 | | | | 60 | 88.0 |
| Th9nL | | | | 75 | 85.0 |
| PM1WT | Basidiomycete PM1 | 5ANH | 496 | 0 | 100 |
| PM13 | | | | 32 | 93.6 |
| PM16 | | | | 45 | 90.9 |
| PM19 | | | | 71 | 85.7 |
| PM19nL | | | | 84 | 83.7 |

**Table S2.** Tv9nL FuncLib design mutations

| ID/position | 162 | 164 | 206 | 265 | 332 | 337 | 393 | 397 | 427 | 455 | 457 | 461 | 464 | # mut |
|---|---|---|---|---|---|---|---|---|---|---|---|---|---|---|
| **Tv9nL** | **F** | **L** | **D** | **F** | **F** | **F** | **A** | **F** | **S** | **I** | **W** | **A** | **A** | **0** |
| Tv9nL.1 | Y | P | Y | T | F | F | F | F | S | L | W | A | A | 6 |
| Tv9nL.2 | F | P | Y | T | Y | F | F | F | S | L | W | A | A | 6 |
| Tv9nL.3 | F | P | Y | T | F | F | F | F | A | L | W | A | S | 7 |
| Tv9nL.4 | F | F | Y | A | F | Y | F | F | S | L | W | A | A | 6 |
| Tv9nL.5 | F | L | Y | A | F | F | F | F | S | V | W | A | A | 4 |
| Tv9nL.6 | F | L | D | F | F | Y | F | L | S | I | W | S | A | 4 |
| Tv9nL.7 | F | F | F | T | F | F | F | L | S | M | W | A | A | 6 |
| Tv9nL.8 | F | I | Y | A | F | Y | A | F | S | L | W | A | A | 5 |
| Tv9nL.9 | F | M | D | M | F | Y | F | F | S | I | W | A | S | 5 |
| Tv9nL.10 | F | M | F | V | F | Y | F | F | S | L | W | A | A | 6 |
| Tv9nL.11 | F | M | Y | A | F | Y | F | F | T | M | W | A | A | 7 |
| Tv9nL.12 | F | M | Y | T | F | F | F | F | T | L | W | A | A | 6 |
| Tv9nL.13 | F | P | D | T | F | Y | A | F | S | I | W | A | A | 3 |
| Tv9nL.14 | F | P | E | M | F | F | F | F | T | I | W | A | A | 5 |
| Tv9nL.15 | F | P | Y | F | F | Y | A | F | S | I | W | A | S | 4 |
| Tv9nL.16 | F | P | Y | A | F | F | S | F | T | L | W | A | S | 7 |
| Tv9nL.17 | F | P | Y | T | F | F | F | F | T | M | W | A | S | 7 |
| Tv9nL.18 | F | P | Y | T | F | F | S | F | T | L | W | A | A | 6 |
| Tv9nL.19 | F | P | Y | T | F | F | V | F | S | L | W | A | A | 5 |
| Tv9nL.20 | F | R | D | I | Y | Y | F | F | S | I | W | A | A | 5 |
| Tv9nL.21 | F | R | E | A | Y | F | F | F | S | M | W | A | A | 6 |
| Tv9nL.22 | I | V | D | V | F | F | F | F | S | I | W | A | S | 5 |
| Tv9nL.23 | Y | L | D | A | F | Y | F | F | T | I | W | A | A | 5 |
| Tv9nL.24 | Y | L | E | F | F | Y | F | F | S | L | W | A | A | 5 |
| Tv9nL.25 | Y | I | D | M | W | F | F | F | S | I | W | A | A | 5 |
| Tv9nL.26 | Y | P | E | A | F | F | F | F | T | I | W | A | A | 6 |
| Tv9nL.27 | Y | P | Y | F | F | Y | A | F | S | I | W | A | A | 4 |
| Tv9nL.28 | Y | P | Y | L | F | F | A | F | S | L | Y | A | A | 6 |
| Tv9nL.29 | Y | R | D | F | F | Y | F | F | S | I | W | A | S | 5 |
| Tv9nL.30 | Y | R | E | F | F | F | F | F | S | L | W | A | A | 5 |

### mut refers to the number of mutations as compared to Tv9nL

**Table S3.** Th3 FuncLib design mutations

| ID/position | 162 | 164 | 206 | 265 | 332 | 337 | 393 | 397 | 427 | 455 | 457 | 461 | 464 | # mut |
|---|---|---|---|---|---|---|---|---|---|---|---|---|---|---|
| **Th3** | **F** | **P** | **N** | **F** | **F** | **F** | **A** | **F** | **S** | **I** | **F** | **A** | **A** | **0** |
| **Th3.1** | F | P | N | **M** | F | F | **S** | F | **N** | I | **W** | A | A | **4** |
| Th3.2 | F | P | F | L | F | Y | T | F | S | V | W | A | A | 6 |
| Th3.3 | F | P | H | F | F | Y | F | F | T | V | W | A | A | 6 |
| Th3.4 | F | P | Y | L | F | Y | T | F | D | V | W | A | A | 7 |
| Th3.5 | F | P | Y | F | F | Y | A | F | T | I | W | A | A | 4 |
| Th3.6 | F | P | N | F | F | Y | A | F | D | I | Y | S | A | 4 |
| **Th3.7** | F | P | **D** | F | F | F | A | F | **D** | **V** | F | A | A | **3** |
| Th3.8 | F | P | D | Y | F | Y | A | F | S | V | W | A | A | 5 |
| Th3.9 | F | P | D | Y | Y | F | A | F | Q | I | W | A | A | 5 |
| **Th3.10** | F | P | **E** | F | F | F | A | F | **D** | I | **Y** | A | A | **3** |
| Th3.11 | F | P | E | L | F | Y | A | F | S | I | W | A | A | 4 |
| Th3.12 | F | P | F | F | F | F | T | F | Q | I | W | A | A | 4 |
| Th3.13 | F | P | F | F | F | Y | A | F | Q | I | W | G | A | 5 |
| **Th3.14** | F | P | **H** | F | F | F | A | F | **N** | I | **W** | A | **S** | **4** |
| Th3.15 | F | P | H | F | F | F | A | F | Q | V | W | A | A | 4 |
| Th3.16 | F | P | H | F | F | F | A | F | T | I | W | G | A | 4 |
| Th3.17 | F | P | R | F | F | Y | T | F | S | I | Y | A | A | 4 |
| Th3.18 | F | P | Y | F | F | F | A | F | Q | V | W | A | S | 5 |
| Th3.19 | F | P | Y | F | F | Y | A | F | Q | V | W | S | A | 6 |
| Th3.20 | F | P | Y | F | F | Y | G | F | Q | I | W | A | A | 5 |
| Th3.21 | F | P | Y | F | F | Y | T | F | Q | V | W | A | A | 6 |
| Th3.22 | F | P | Y | F | Y | Y | A | F | Q | I | W | A | A | 5 |
| Th3.23 | F | P | Y | Y | F | F | A | F | Q | V | W | A | A | 5 |
| Th3.24 | F | P | Y | Y | F | Y | A | F | Q | V | Y | A | A | 6 |
| Th3.25 | Y | P | N | F | F | Y | A | F | A | V | F | A | A | 4 |
| Th3.26 | Y | P | N | F | F | Y | T | F | T | I | Y | A | A | 5 |
| Th3.27 | Y | P | F | F | F | F | T | F | S | I | W | A | A | 4 |
| Th3.28 | Y | P | F | F | F | Y | G | F | T | I | W | A | A | 6 |
| Th3.29 | Y | P | Y | F | F | F | G | F | S | V | W | A | A | 5 |
| Th3.30 | Y | P | Y | F | F | Y | A | F | A | V | W | A | A | 6 |

### mut refers to the number of mutations as compared to Th3

**Table S4.** Kinetic parameters of FuncLib designs with various substrates

| Enz\Sub | Kinetic constants | ABTS | DMP | GUA | SA | VLA |
|---|---|---|---|---|---|---|
| Th3 | $K_M$ (µM)<br>$k_{cat}$ ($sec^{-1}$)<br>$k_{cat}$ / $K_M$ ($sec^{-1}$ $mM^{-1}$) | 21.8 ± 0.9<br>1.88 ± 0.02<br>86.2 ± 3.7 | 1280 ± 120<br>0.520 ± 0.021<br>0.406 ± 0.041 | 1790 ± 160<br>0.347 ± 0.012<br>0.194 ± 0.019 | n.d. (very low efficiency) | 11.6 ± 1.6<br>2.77 ± 0.14<br>0.239 ± 0.035 |
| Th3.1 | $K_M$ (µM)<br>$k_{cat}$ ($sec^{-1}$)<br>$k_{cat}$ / $K_M$ ($sec^{-1}$ $mM^{-1}$) | 16.3 ± 0.8<br>0.859 ± 0.013<br>52.7 ± 2.7 | 1030 ± 80<br>0.611 ± 0.024<br>0.593 ± 0.052 | 555 ± 85<br>0.099 ± 0.006<br>0.178 ± 0.029 | n.d. (very low efficiency) | 32.0 ± 5.4<br>1.93 ± 0.17<br>0.060 ± 0.011 |
| Th3.7 | $K_M$ (µM)<br>$k_{cat}$ ($sec^{-1}$)<br>$k_{cat}$ / $K_M$ ($sec^{-1}$ $mM^{-1}$) | 65.6 ± 5.5<br>15.6 ± 0.6<br>237 ± 22 | 225 ± 3<br>13.5 ± 0.1<br>60.1 ± 0.8 | 1320 ± 70<br>3.98 ± 0.06<br>3.02 ± 0.17 | 300 ± 15<br>10.5 ± 0.2<br>35.1 ± 1.9 | 38.4 ± 2.3<br>8.94 ± 0.30<br>0.233 ± 0.016 |
| Th3.10 | $K_M$ (µM)<br>$k_{cat}$ ($sec^{-1}$)<br>$k_{cat}$ / $K_M$ ($sec^{-1}$ $mM^{-1}$) | 9.17 ± 0.96<br>3.36 ± 0.09<br>366 ± 40 | 17.1 ± 1.0<br>4.10 ± 0.05<br>240 ± 14 | 667 ± 36<br>1.55 ± 0.03<br>2.32 ± 0.13 | 152 ± 14<br>4.41 ± 0.13<br>29.0 ± 2.8 | 20.4 ± 2.0<br>7.85 ± 0.36<br>0.385 ± 0.042 |
| Th3.14 | $K_M$ (µM)<br>$k_{cat}$ ($sec^{-1}$)<br>$k_{cat}$ / $K_M$ ($sec^{-1}$ $mM^{-1}$) | 8.55 ± 0.65<br>7.83 ± 0.16<br>916 ± 72 | 44.0 ± 3.2<br>1.84 ±0.03<br>41.8 ± 3.1 | 771 ± 43<br>1.00 ± 0.02<br>1.30 ± 0.08 | 185 ± 13<br>2.45 ± 0.08<br>13.2 ± 1.0 | 10.5 ± 1.6<br>5.45 ± 0.36<br>0.519 ± 0.086 |

**Supplementary Figures**

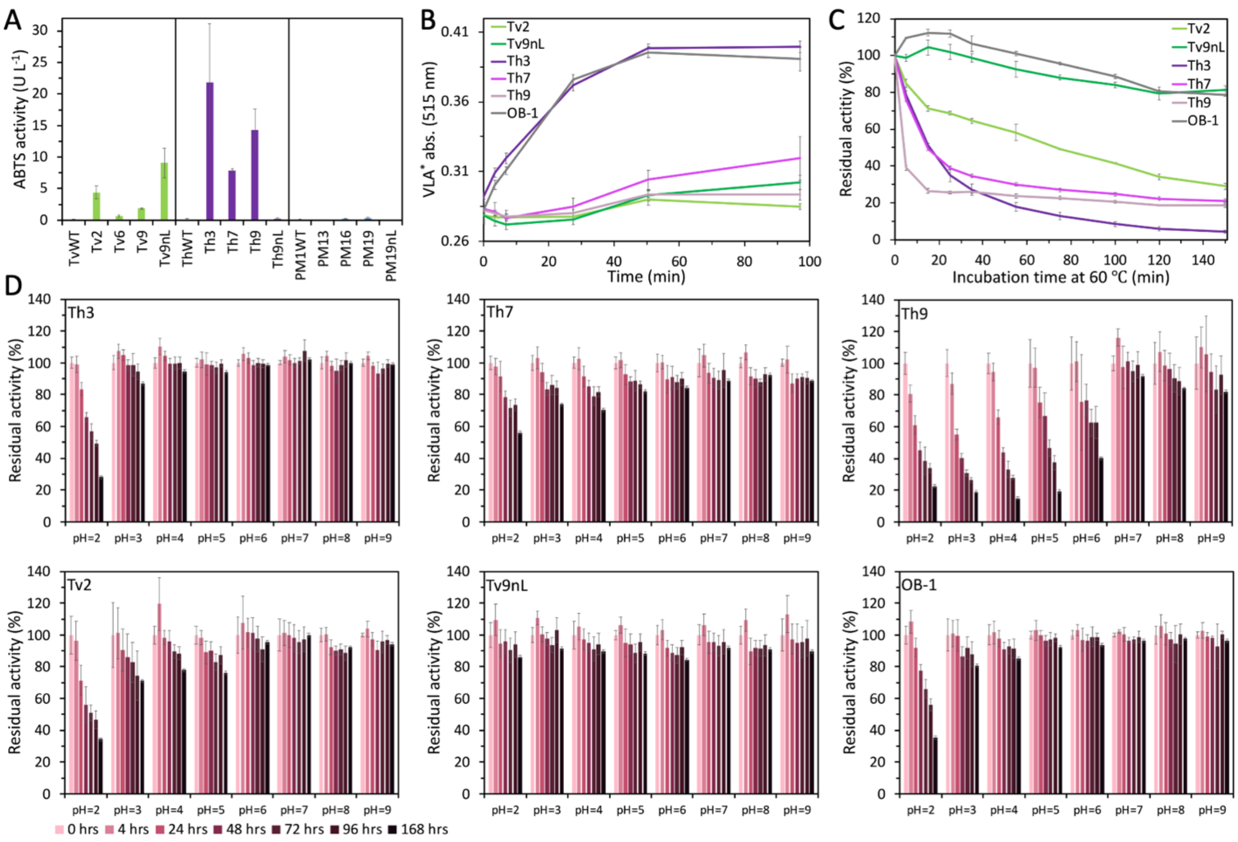


**Figure S1.** Stability and activity of PROSS designs. (A) Screening of HRPLs from *Trametes versicolor* (Tv), *Trametes hirsuta* (Th) and basidiomycete PM1 (PM1). Activity of PROSS designs against ABTS indicates that three Tv and Th designs are functionally expressed while their wildtype progenitors are not. The results are presented as the mean ± S.D. of three independent biological replicates. 1U is defined as 1 μmol/min. (B) Oxidation of the high-redox potential mediator VLA (to form VLA*) was measured by incubating yeast supernatants of best PROSS designs with 20 mM VLA at pH=4.0 and measuring the absorption of VLA* at 515 nm. (C) Kinetic thermostability ($t_{1/2}$) profiles were determined by incubating the yeast supernatants of best PROSS designs at 60 °C and measuring the residual activity at times 0-150 minutes, compared to the initial activity. (D) pH stability profiles were determined by incubating the yeast supernatants of the active PROSS designs at 100 mM borate-citrate-phosphate buffer with pH values ranging from 2 to 9 and measuring the residual activity at times 0-168 hours, compared to the initial activity at each pH. (B-D) results are presented as the mean ± S.D. of three independent experiments.

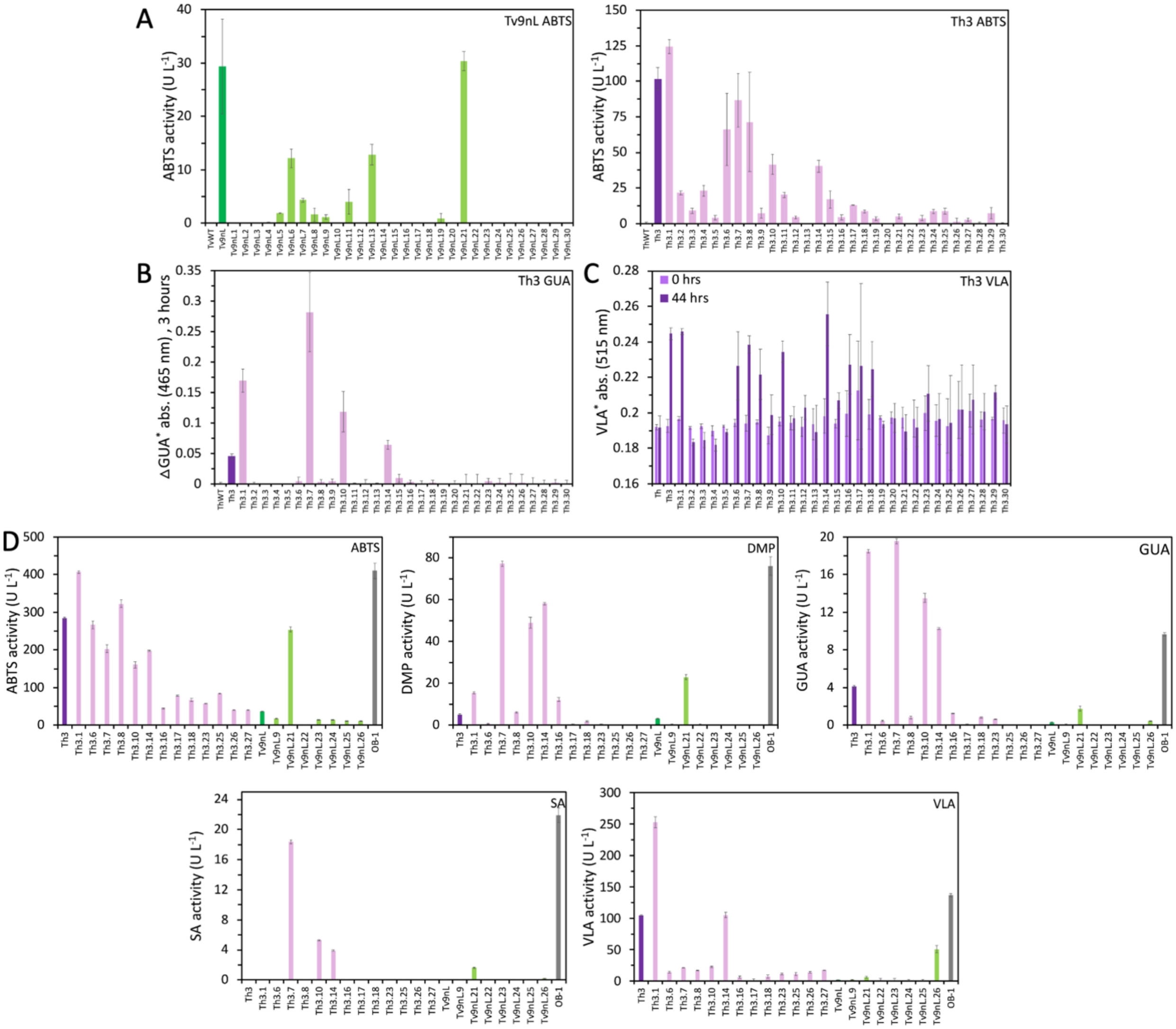


**Figure S2.** Diverse selectivity profiles of Tv9nL and Th3 FuncLib designs. (A-C) Initial screening of all FuncLib designs expressed under restrictive growth conditions in 96-well plates. Activity of yeast supernatants was measured at pH=4.0 against ABTS for (A) Tv9nL designs and Th3 designs, and against (B) GUA and (C) VLA for Th3; activity against GUA and VLA was measured for Tv9nL designs as well but indicated for no significant oxidation (data not shown). The results are presented as the mean ± S.D. of three independent biological replicates. (D) Second screening of selected Tv9nL and Th3 FuncLib designs expressed in rich expression media. Activity of yeast supernatants was measured against five substrates (indicate top-right) at pH=4.0. The results are presented as the mean ± S.D. of three independent experiments.

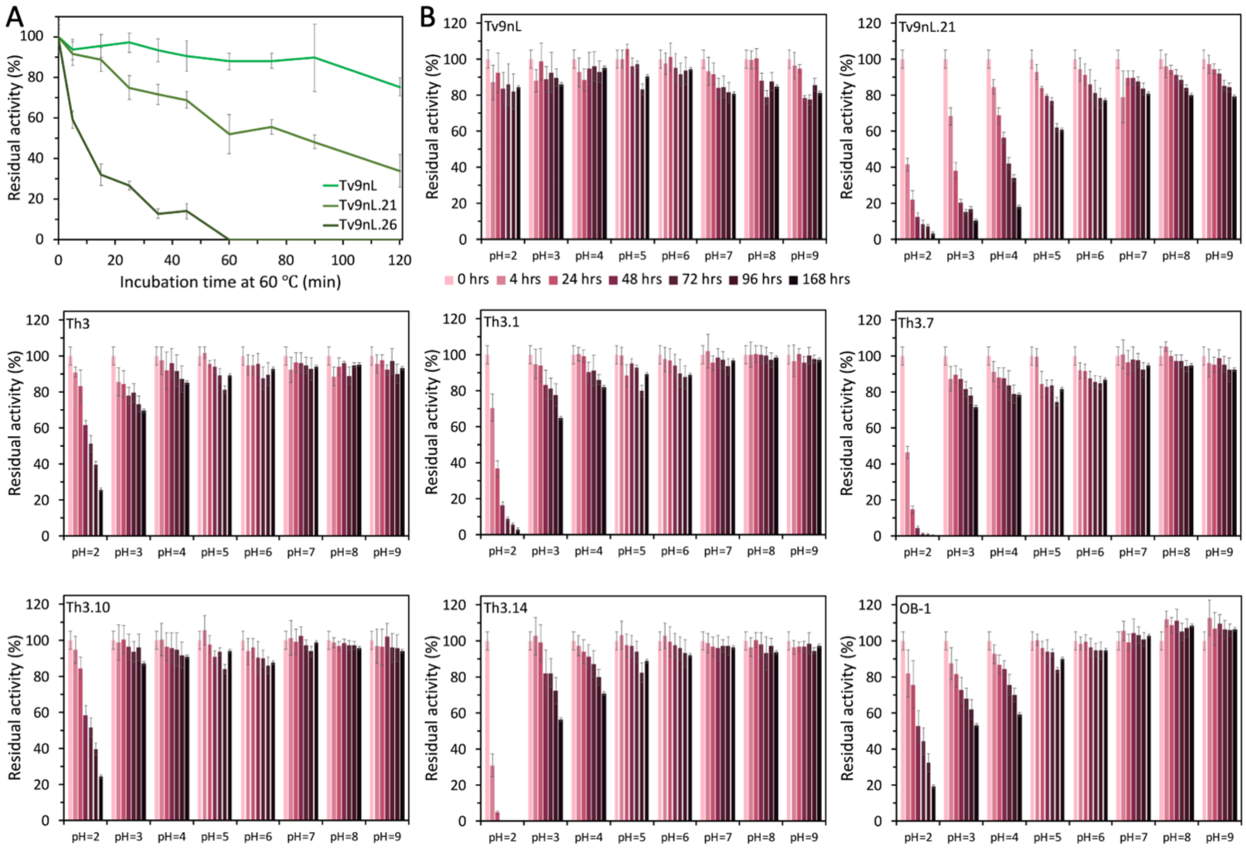


**Figure S3.** Stability of FuncLib designs. (A) Kinetic thermostability ($t_{1/2}$) profiles were determined by incubating the yeast supernatants of selected Tv9nL FuncLib designs at 60 °C and measuring the residual activity at times 0-120 minutes, compared to the initial activity. (D) pH stability profiles were determined by incubating the yeast supernatants of selected FuncLib designs at 100 mM borate-citrate-phosphate buffer with pH values ranging from 2 to 9 and measuring the residual activity at times 0-168 hours, compared to the initial activity at each pH. All of the results are presented as the mean ± S.D. of three independent experiments.

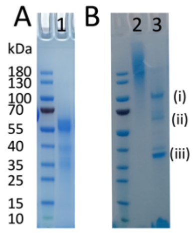


**Figure S4.** SDS-PAGE analysis of purified Th3. (A) Purified Th3 after ammonium-sulfate precipitation, anion-exchange chromatography and size exclusion chromatography (lane 1). (B) Purified Th3 after 100% ammonium-sulfate precipitation and hydrophobic interaction chromatography (lane2). Hyper-glycosylated fraction (lane 2) was treated with the N-glycosidase PNGaseF, and the three major bands (i-iii; lane 3) were analyzed by proteolytic mass-spectrometry.

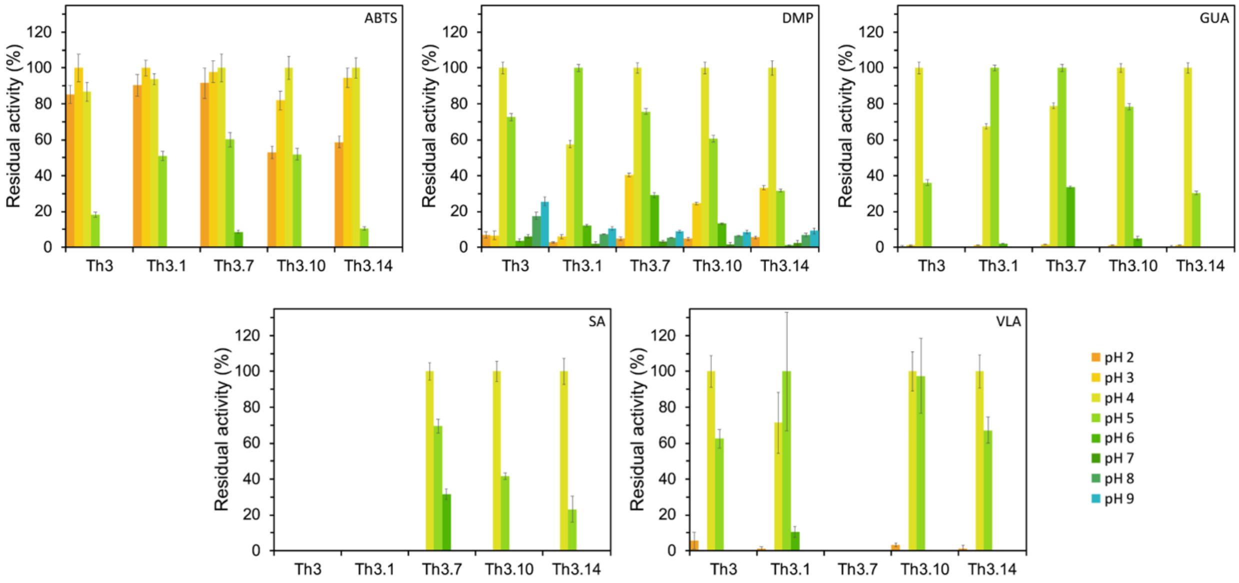


**Figure S5.** pH-dependent activity of FuncLib designs. The pH-dependent activity profiles were determined by measuring the yeast supernatant activity of selected FuncLib designs with the substrates (indicate top-right) at a range of pHs (100 mM borate-citrate-phosphate buffer with pH values ranging from 2 to 9). The activities were normalized to the activity at optimal pH for each protein-substrate pair. All of the results are presented as the mean ± S.D. of three independent experiments.
